# The ICF syndrome protein CDCA7 harbors a unique DNA-binding domain that recognizes a CpG dyad in the context of a non-B DNA

**DOI:** 10.1101/2023.12.15.571946

**Authors:** Swanand Hardikar, Ren Ren, Zhengzhou Ying, John R. Horton, Matthew D. Bramble, Bin Liu, Yue Lu, Bigang Liu, Jiameng Dan, Xing Zhang, Xiaodong Cheng, Taiping Chen

**Affiliations:** Department of Epigenetics and Molecular Carcinogenesis, The University of Texas MD Anderson Cancer Center, Houston, Texas 77030, USA; Program in Genetics and Epigenetics, The University of Texas MD Anderson Cancer Center UTHealth Graduate School of Biomedical Sciences, Houston, Texas 77030, USA

## Abstract

*CDCA7*, encoding a protein with a C-terminal cysteine-rich domain (CRD), is mutated in immunodeficiency, centromeric instability and facial anomalies (ICF) syndrome, a disease related to hypomethylation of juxtacentromeric satellite DNA. How CDCA7 directs DNA methylation to juxtacentromeric regions is unknown. Here, we show that the CDCA7 CRD adopts a unique zinc-binding structure that recognizes a CpG dyad in a non-B DNA formed by two sequence motifs. CDCA7, but not ICF mutants, preferentially binds the non-B DNA with strand-specific CpG hemi-methylation. The unmethylated sequence motif is highly enriched at centromeres of human chromosomes, whereas the methylated motif is distributed throughout the genome. At S phase, CDCA7, but not ICF mutants, is concentrated in constitutive heterochromatin foci, and the formation of such foci can be inhibited by exogenous hemi-methylated non-B DNA bound by the CRD. Binding of the non-B DNA formed in juxtacentromeric regions during DNA replication provides a mechanism by which CDCA7 controls the specificity of DNA methylation.

## Introduction

The predominant conformation of DNA in cells is the canonical right-handed B-form double-stranded helix. However, a variety of non-canonical DNA conformations, such as hairpins, cruciforms, left-handed double-helical Z-DNA, triple-stranded H-DNA, G-quadruplexes, and RNA-like four-way junction, have also been recognized (1–8). Non-B-form structures can affect DNA-dependent processes, including transcription, replication, recombination and repair, and have been implicated in mutagenesis and genetic instability that are associated with various diseases (1,9,10). Recent evidence suggests that non-B-form DNA is particularly enriched at centromeres in eukaryotes (11–13), which has led to the hypothesis that non-B DNA structures contribute to centromere specification (11). Presumably, some of the biological effects of non-B DNA conformations are mediated by proteins that recognize and/or stabilize them. Indeed, numerous non-B DNA-binding proteins have been reported (14–16). However, the molecular mechanisms by which these proteins interact with their corresponding non-B DNA substrates, as well as the functional significance of most such interactions, have not been well characterized.

Immunodeficiency, centromeric instability and facial anomalies (ICF) syndrome is a rare autosomal recessive disease characterized by immunoglobulin deficiency, facial dysmorphism, intellectual disability, developmental delay, and genomic instability involving the juxtacentromeric regions of chromosomes 1, 9 and 16, (17,18). These chromosomes have relatively large centromeric and pericentromeric (hereafter peri/centromeric) regions. ICF patients usually suffer from recurrent infections in early childhood (19–21).

A hallmark of ICF syndrome is hypomethylation of specific genomic regions, most notably classical satellite repeats in peri/centromeric regions (22). Four ICF-related genes have been identified – *DNMT3B* (DNA methyltransferase 3B), *ZBTB24* (zinc finger- and BTB domain-containing 24), *CDCA7* (cell division cycle associated 7), and *HELLS* (helicase, lymphoid-specific, also known as lymphoid-specific helicase, *LSH*) – with cases carrying different gene mutations being designated, respectively, as ICF1 (OMIM #242860, *DNMT3B*), ICF2 (OMIM #614069, *ZBTB24*), ICF3 (OMIM #616910, *CDCA7*), ICF4 (OMIM #616911, *HELLS*), and ICFX (unknown) (23–27). DNMT3B is a *de novo* DNA methyltransferase that establishes DNA methylation patterns during development (24). HELLS, a DNA helicase involved in chromatin remodeling, regulates both *de novo* and maintenance DNA methylation in an ATPase-dependent manner (28–31). Recent studies suggest that ZBTB24 and CDCA7 function upstream of HELLS in a molecular pathway that regulates DNA methylation. Specifically, ZBTB24, a C2H2-zinc finger transcription factor, induces *CDCA7* transcription, and CDCA7 recruits HELLS to centromeric satellite repeats, among other regions, to facilitate DNA methylation (32–37). Thus, CDCA7 plays a key role in determining the specificity of the ZBTB24-CDCA7-HELLS axis in the regulation of DNA methylation.

*CDCA7* was identified as a c-Myc-responsive gene (38). It is periodically expressed in the cell cycle and reaches the highest level between G1 and S phase and has been implicated in transcriptional regulation, tumorigenesis, and hematopoietic stem cell emergence (39–44). However, the fundamental functions of CDCA7 remain poorly understood. In addition to several small functional regions, i.e., a leucine zipper motif, a c-Myc/14-3-3 interaction motif, and a nuclear localization signal, CDCA7 harbors a C-terminal cysteine-rich domain (CRD) including four copies of CXXC motif (27,41). Of note, the identified ICF3 missense mutations in CDCA7 alter three residues in the CRD (27), highlighting the functional importance of the domain. Previous studies have shown that ICF3 mutations disrupt the recruitment of the CDCA7-HELLS complex to chromatin (33,36). Nevertheless, the precise role of the CDCA7 CRD and the chromatin component and feature being recognized are unknown.

In this study, we provide structural and DNA-binding data demonstrating that the CDCA7 CRD adopts a unique three-zinc-containing structure that binds a non-B-form DNA in a CpG-specific manner. The three ICF3 missense mutations abolish binding by disrupting, respectively, the interactions with a Zn^2+^, a guanine of a CpG dinucleotide, and a phosphate group of the DNA backbone. The non-B DNA is formed by two sequence motifs. A 13-mer motif 1 is present throughout the human genome, and an

11-mer motif 2 is highly enriched in centromeric alpha satellite (αSat) repeats. Strand-specific CpG methylation in the non-B DNA exhibits the opposite effects on CDCA7 binding – positively regulated by motif 1 methylation and negatively regulated by motif 2 methylation. We also show that wild-type (WT), but not ICF3-mutant, CDCA7 is concentrated in constitutive heterochromatin foci during the S phase of the cell cycle. The hemi-methylated non-B DNA preferentially bound by the CDCA7 CRD, when introduced in cells, can inhibit CDCA7 foci formation and induce hypomethylation of centromeric satellite DNA. We propose that CDCA7 recruits HELLS and the DNA methylation machinery to centromeric regions by recognizing the non-B DNA formed during DNA replication.

## Results

### Wild-type CDCA7, but not ICF3 mutants, is concentrated in constitutive heterochromatin foci at S phase

Human (h) and mouse (m) CDCA7 are highly conserved, with ∼97% sequence identity in the CRD, where the ICF3 missense mutations are located (Fig. 1A). We previously showed that CDCA7 recruits HELLS to heterochromatin to facilitate DNA methylation of the centromeric minor satellite repeats in mouse embryonic stem cells (mESCs) (37). Mutagenesis analysis indicated that the leucine zipper motif is required for CDCA7 to interact with HELLS, whereas an ICF3 missense mutation and even deletion of the entire CRD showed no effect on CDCA7-HELLS interaction (Fig. S1). Our results were consistent with previous observation that ICF3 missense mutations do not affect the formation of the CDCA7-HELLS complex but prevent the recruitment of the complex to chromatin (33,36).

**Fig. 1.**
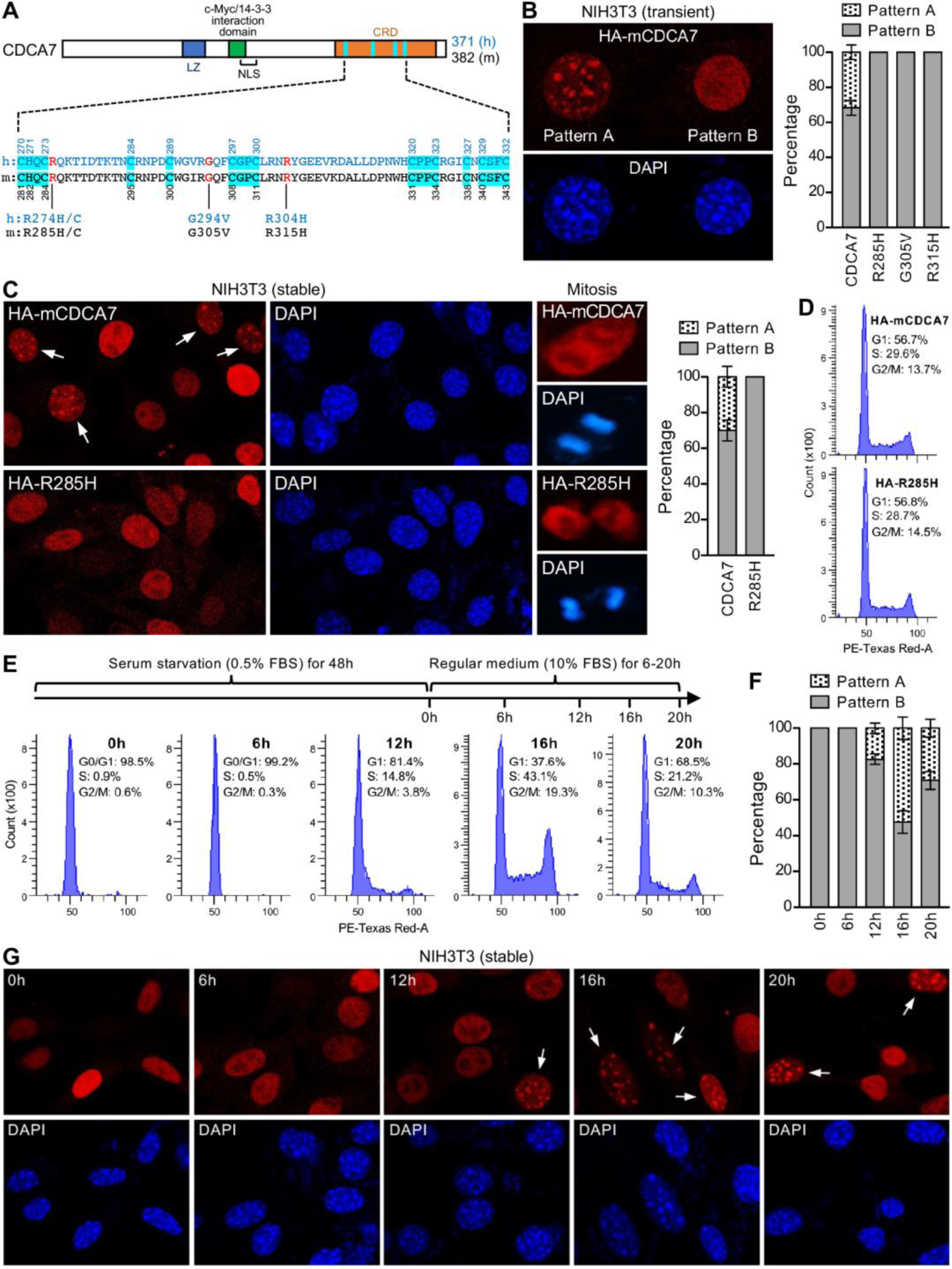
CDCA7, but not ICF3 mutants, is concentrated in heterochromatic foci at S phase. (**A**) CDCA7 protein with the defined domains: a leucine zipper (LZ) motif, a c-Myc/14-3-3 interaction motif, a nuclear localization signal (NLS), and a cysteine-rich domain (CRD) that contains four CXXC motifs (highlighted in Cyan). The amino acid sequences from the 1^st^ to 4^th^ CXXC motifs in human (h) and mouse (m) CDCA7 are shown, with the Cys and His residues involved in zinc coordination being highlighted in Cyan and numbered. The ICF3 missense mutations are indicated. (**B**) IF data showing that HA-tagged mCDCA7 transiently expressed in NIH3T3 cells exhibits two nuclear localization patterns, pattern A (enrichment in heterochromatin foci) and pattern B (no enrichment in heterochromatin foci), whereas ICF3 mutant proteins (R285H, G305V and R315H) exhibit only pattern B. Shown are representative images (left) and percentages (right) of the two patterns. (**C**) IF data with stable expression of HA-tagged mCDCA7 or the R285H mutant in NIH3T3 cells. Shown are representative images (left) and percentages (right) of the two localization patterns. During mitosis, both WT and mutant mCDCA7 proteins are localized throughout the cell, excluding the chromosomes. (**D**) Flow cytometry analysis showing that NIH3T3 cells stably expressing WT or mutant mCDCA7 have similar cell cycle profiles. (**E-G**) Cell cycle synchronization experiments using serum starvation, which suggest mCDCA7 enrichment in heterochromatin foci during S phase. Shown are the cell cycle profiles (**E**), percentages of the two localization patterns (**F**), and representative images (**G**) at different timepoints. The quantification data in (**B, C** and **F**) represent mean ± SD from three experiments, with >100 interphase cells being counted each time. The arrows in (**C** and **G**) indicate pattern A cells.

To gain insights into the effect of ICF3 mutations on CDCA7 chromatin association, we examined CDCA7 cellular localization. Immunofluorescence (IF) analysis revealed that HA-tagged WT mCDCA7 and the R285H, G305V and R315H mutants (equivalent to the R274H, G294V and R304H ICF3 mutants in hCDCA7; Fig. 1A) all localized in the nuclei in transiently transfected NIH3T3 cells (Fig. 1B). WT mCDCA7 was highly enriched in constitutive heterochromatin foci (pattern A) – marked by DAPI bright spots – in a considerable fraction (∼30%) of transfected cells, although the majority (∼70%) of cells showed no such enrichment (pattern B). In contrast, the ICF3 mutant proteins failed to be concentrated in heterochromatin foci and exhibited only the diffused pattern B (right panel of Fig. 1B). Similar results were obtained when HA-tagged hCDCA7 proteins were expressed in the human cervical cancer cell line HeLa: concentration of WT hCDCA7, but not the R274H ICF3 mutant, in constitutive heterochromatin foci, as labeled by the histone H3 lysine 9 trimethyl (H3K9me3) mark (Fig. S2).

We also established stable NIH3T3 cell lines expressing HA-tagged mCDCA7 or the R285H mutant (Fig. 1C). In agreement with the results of transient transfection (Fig. 1B), IF analysis of the stable cell lines showed that WT mCDCA7 exhibited both pattern A and pattern B, accounting for ∼30% and ∼70% of the interphase cells, respectively, but the R285H mutant exhibited pattern B in all interphase cells (Fig. 1C). During mitosis, both WT and mutant mCDCA7 proteins were present throughout the cells, excluding the chromosomes (Fig. 1C, images on the right). The two stable cell lines (WT and R285H mutant) showed no difference in viability and proliferation, and flow cytometry analysis revealed similar cell cycle profiles (Fig. 1D).

The presence of CDCA7 nuclear foci only in a fraction of cells raises the possibility of CDCA7 localization patterns being regulated during the cell cycle. Thus, we first synchronized HA-mCDCA7-expressing NIH3T3 cells at the G0/G1 phase by serum starvation (0.5% FBS) for 48h, followed by culturing them in regular medium (containing 10% FBS) for 6, 12, 16 and 20h, respectively. As revealed by cell cycle analysis, almost all cells were arrested at G0/G1 phase initially (0h timepoint) and remained at G0/G1 phase at the 6h timepoint, small fractions reached S (∼15%) and G2/M (∼4%) phases at the 12h timepoint, substantially larger fractions were at S(∼43%) and G2/M (∼19%) phases at the 16h timepoint, and most cells apparently had entered the next G1 phase at the 20h timepoint (Fig. 1E). CDCA7 nuclear foci (pattern A cells) were not observed at 0h and 6h but appeared in ∼17%, ∼52% and ∼29% of the cells at 12h, 16h and 20h, respectively (Fig. 1, F and G). We conclude that pattern A cells are mostly at S phase, although the localization pattern may persist to G2 phase in some cells. Taken together, our results indicate that CDCA7 is enriched in constitutive heterochromatin during DNA replication and that the ICF3 missense mutations in the CRD disrupt such enrichment.

### The CDCA7 CRD binds a specific non-B DNA

Our finding suggests that determining how the CRD targets CDCA7 to constitutive heterochromatin is key to understanding the mechanism by which the ZBTB24-CDCA7-HELLS axis specifically regulates methylation of satellite DNA repeats in juxtacentromeric regions (37). The CDCA7 CRD has been implicated in DNA binding (33). However, electrophoretic mobility shift assay (EMSA) showed that the CDCA7 CRD failed to bind DNA sequences containing the repeating units of several common satellite DNAs in the peri/centromeric regions of human and mouse genomes (Fig. S3A).

One possibility is that the CDCA7 CRD recognizes a specific DNA motif or structure that is present in heterochromatic regions of both human and mouse cells. To explore the possibility, we performed <u>S</u>ystematic <u>E</u>volution of <u>L</u>igands by <u>EX</u>ponential Enrichment (SELEX) (Fig. S3B), a technique for identifying single-stranded (ss) ‘aptamers’ recognized by sequence-specific binding proteins (45). By screening a library of 30-mer random ssDNA sequences using a recombinant GST fusion protein comprising mCDCA7 CRD, we identified two highly similar sequences, named Seq-1 and Seq-2 (Fig. 2A). EMSA verified the binding of both sequences by the CRD of mCDCA7 and hCDCA7 and the disruption of binding by ICF3 missense mutations (Fig. 2, B and C, F probe = Seq-1; Fig. S4, ss-2 = Seq-2). The binding was highly specific, as the CDCA7 CRD failed to bind the reverse complementary probe (R probe), a double-stranded (ds) DNA probe (annealed F/R probes) (Fig. 2B), an RNA probe with the same sequence as F probe, and an RNA/DNA hybrid probe (annealed RNA F probe and DNA R probe) (Fig. 2D).

**Fig. 2.**
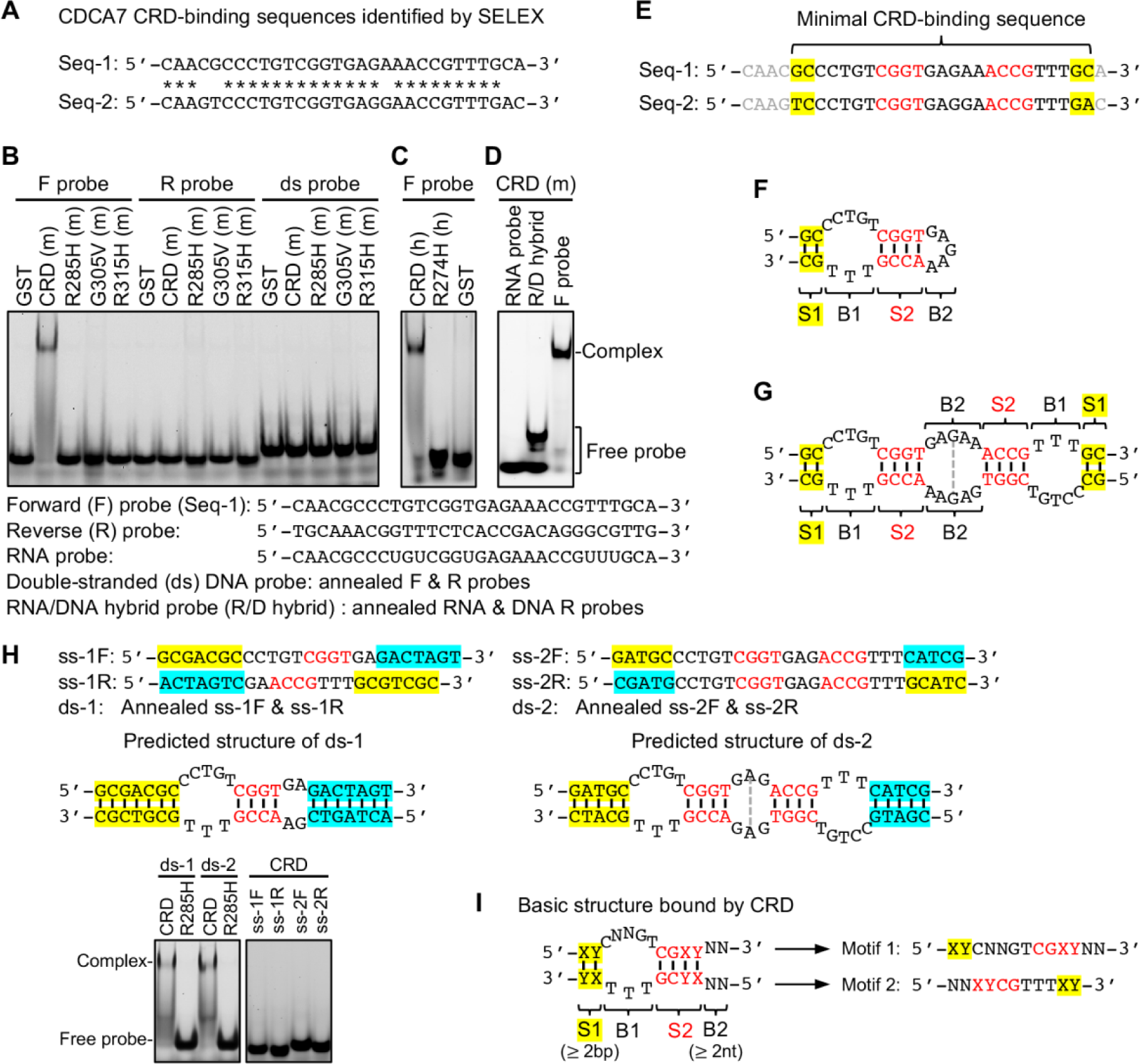
The CDCA7 CRD binds a specific non-B DNA. (**A**) The two sequences, Seq-1 and Seq-2, identified by SELEX (identical bases indicated by *). (**B-D**) EMSA experiments using GST fusion proteins comprising of the CRD of mCDCA7, hCDCA7 or CRD with ICF3 mutations. The probes used for EMSA were shown below: ssDNA probes (F probe: identical to Seq-1; R probe: reverse complementary to F probe), RNA probe (sequence identical to that of F probe), dsDNA probe (annealed F and R probes), and RNA/DNA (R/D) hybrid probe (annealed RNA and DNA R probes). (**E**) The minimal CRD-binding sequence determined by mutagenesis (see Fig. S4). (**F** and **G**) Possible non-B structures formed by the CRD-binding sequence of Seq-1, through intra-strand (**F**) or inter-strand (**G**) base pairing. (**H**) EMSA data with ds probes by annealing ss oligos, which showed that the non-B structures shown in (**F** and **G**) can both be bound by mCDCA7 CRD. Note that the ssDNAs (ss-1F, ss-1R, ss-2F and ss-2R) used to form ds probes are not bound by the CRD. (**I**) The basic structure bound by the CDCA7 CRD, as defined by mutagenesis results shown in figs. S4-S6, and the two sequence motifs, motif 1 and motif 2, that form the basic structure. N means any unpaired nucleotide, and X:Y means any base pair.

By testing different fragments in the 30-mer Seq-1, we determined that the four nucleotides at the beginning (5’ end) and one nucleotide at the end (3’ end) are not required for binding by mCDCA7 CRD (Fig. S4, ss-3 to ss-13). The 25-mer CRD-binding sequences of Seq-1 and Seq-2 (Fig. 2E) contain two short dyad symmetries, a pair of tetranucleotides (CGGT and ACCG) in the middle (highlighted in red) and a pair of dinucleotides (GC and GC in Seq-1, TC and GA in Seq-2) at the ends (highlighted in yellow). Two possible non-B DNA structures could be formed: a hairpin by intra-strand base pairing, with two stems (S1 and S2) and two bubbles (B1 and B2) (Fig. 2F) or a more complex structure by inter-strand base pairing, with two symmetric halves (Fig. 2G). Using dsDNA probes by annealing single-stranded oligos, we found that both the hairpin (Fig. 2F) and more complex structure (Fig. 2G) can be bound by mCDCA7 CRD (Fig. 2H). Binding of the non-B DNA was also verified with various other probes (Fig. S4, ss-14, ss-15 and ss-18).

Extensive mutagenesis of the non-B DNA revealed the following requirements for CDCA7 binding. First, stem S1 must have at least 2 bps immediately next to bubble B1, and the sequence is not important (Fig. S4, ss-6 to ss-13; Fig. S6, ss-49 to ss-51, ss-121 to ss-125). Second, the number of nucleotides that form bubble B1 (CCTGT and TTT) cannot be changed (Fig. S5, ss-20 to ss-45; Fig. S6, ss-52 to ss-54, ss-118 to ss-120), and the TTT sequence is essential (Fig. S6, ss-109 to ss-117), but the CCTGT sequence can be CNNGT (N means any nucleotide) (Fig. S6, ss-55 to ss-69). Third, stem S2 must be 4 bps (Fig. S4, ss-17; Fig. S6, ss-70 to ss-81, ss-97 to ss-108), and the C:G and G:C base pairs at positions 1 and 2 are essential, whereas the base pairs at positions 3 and 4 can be formed by any nucleotides (Fig. S6, ss-126 to ss-137). Fourth, bubble B2 must have at least two nucleotides on each strand (Fig. S5, ss-46 to ss-48), but the sequence does not seem to be important (Fig. S6, ss-82 to ss-96). While DNA binding was examined by EMSA, the binding affinities of some probes were measured by isothermal titration calorimetry (ITC) using the CRD of mCDCD7 or hCDCA7, which confirmed the EMSA results (figs. S7 and S8).

In summary, the basic structure bound by the CDCA7 CRD is a non-B-form DNA with a stem S1 of at least 2 bps (shown as X:Y and Y:X, meaning any base pairs), a bubble B1 formed by CNNGT on one strand and TTT on the other, a 4-bp stem S2 containing an essential CpG dyad, followed by a bubble B2 formed by at least 2 mismatches (Fig. 2I). The two sequence elements that form the basic non-B structure are referred to as motif 1 (5’-<u>XY</u>CNNGT<u>CGXY</u>NN-3’) and motif 2 (5’-NN<u>XYCG</u>TTT<u>XY</u>-3’) with base pairs underlined (Fig. 2I).

### Structural investigations

We used a set of oligonucleotides for co-crystallization trials, with a length varying from 36-nt to 26-nt by reducing one nucleotide at a time from both ends. The 24-nt is a minimal length for mCDCA7 CRD to bind, as further shortened 22-nt and 20-nt resulted in >20X and >70X reduced binding affinity, respectively (Fig. 3A, Fig. S7A). We determined five structures of mCDCA7 CRD in complex with 36-nt (in two different crystallographic cell dimensions), 34-nt, 32-nt and 26-nt in the resolution range of 2.6 Å to 1.6 Å (Fig. S9 and Table S1). The protein-DNA complexes were all crystallized in space group *C*2, with varied cell dimensions of crystal lattices, containing either one or two protein-ssDNA complexes per crystallographic asymmetric unit. In addition, we determined mCDCA7 or hCDCA7 CRD in complex with a hemi-methylated CpG site in the context of 32-nt (see below). Thus, we describe the structures of complex with the 32-nt determined at 1.9 Å resolution. All 32 nucleotides are clearly resolved in the electron densities.

**Fig. 3.**
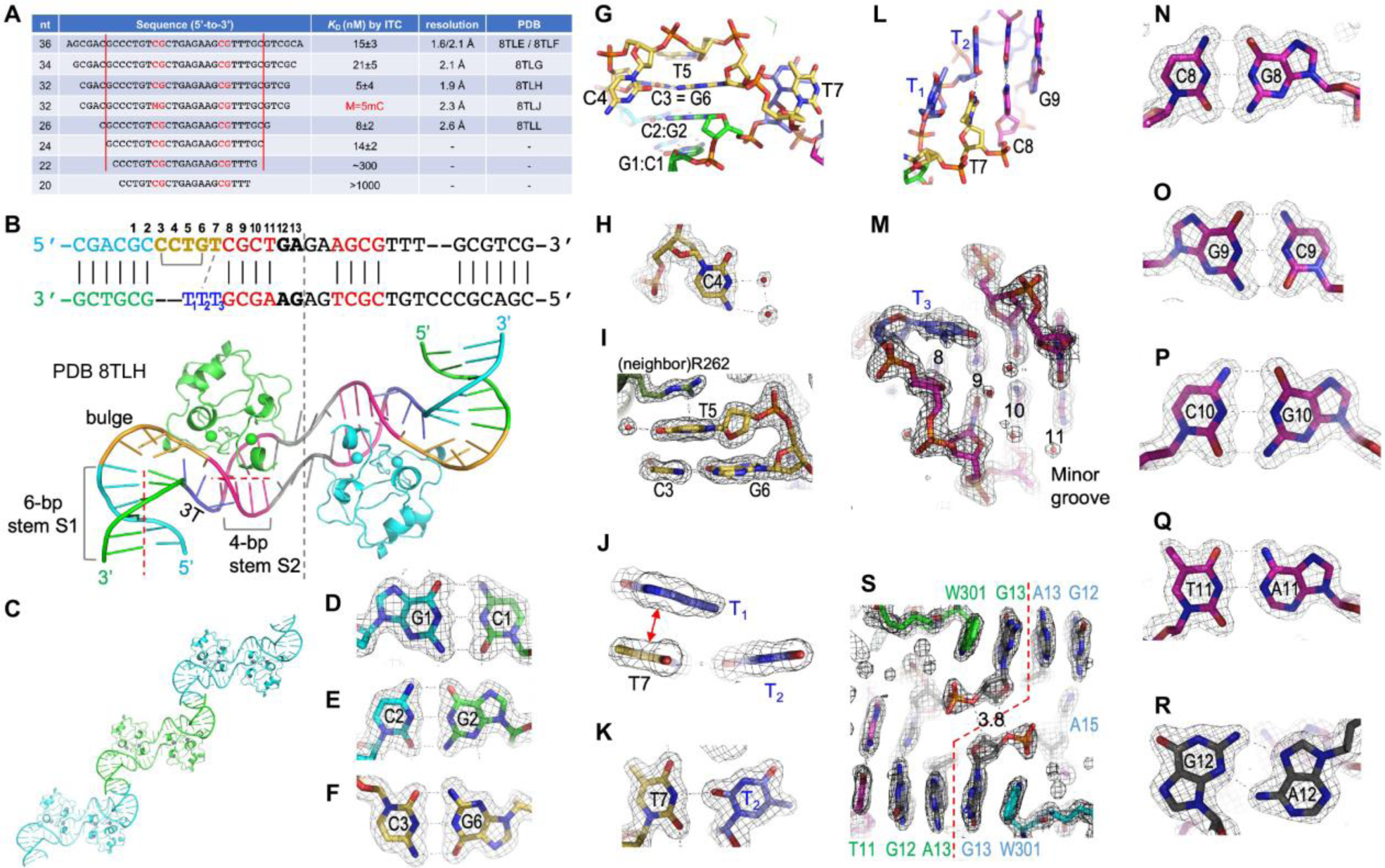
The CDCA7-bound DNA adopts a non-B-form conformation. (**A**) Summary of DNA sequences used for ITC binding measurements and structural information (PDB accession number and corresponding resolutions). (**B**) The 32-nt oligos used for co-crystallization and the nucleotide numbering above the sequence. The subscribed T_1_, T_2_ and T_3_ are used for the triple-T element of the bottom strand. The 6-bp stem S1, the 4-bp stem S2, bulged region and 3T are labeled and colored coded. The two dashed red lines indicate the axes of the two stems, which are in a ∼90° turn. (**C**) Two neighboring CDCA7 CRD-DNA complexes form a pseudo-continuous duplex between the two DNA stems. (**D**) G1:C1 base pair. (**E**) C2:G2 base pair. (**F** and **G**) An intra-strand C3 and G6 base pair (**F**), which stacks with the last base pair of stem S1 (**G**). (**H**) C4 protrudes from the bulge and is unstacked. (**I**) T5 stacks with C3:G6 on one side and Arg362 of neighboring molecule on the other side. (**J** and **K**) T7 stacks with T_1_ and H-bonds with T_2_. (**L**) T7 and T_2_ mismatch stacks with the first C:G base pair of stem S2. (**M**) T_3_ is located in the minor groove of CpG dinucleotide at positions 8 and 9. (**N-Q**) The four base pairs of stem S2. (**R**) A purine mismatch at position 12. (**S**) Trp301 stacks with an extrahelical G13. The dashed red line indicates the two symmetric halves of the crystalized complexes. Two pairs of deoxyribose C4’ atom of G13 and the O3’ atom of G13 from the opposite strand are in close contact.

### Non-B-form DNA

Instead of forming a hairpin structure, the two annealed single-stranded oligos couple to each other and form a non-B DNA conformation with two symmetric halves, and each half is bound by one CDCA7 CRD (Fig. 3B). This implies that in the crystal the protein domain and ssDNA is in equal molar ratio. As expected, the non-B DNA can be divided into four sections: a 6-bp stem S1, a bubble formed by a 5-nt bulged loop of the top strand and a 3-nt triple T of the bottom strand, a 4-bp stem S2 and two purine mismatches. The 6-bp stem S1 was coaxially stacked with the neighboring DNA molecule, forming a pseudo-continuous duplex between the two DNA molecules throughout the crystal lattice (Fig. 3C). We numbered the DNA sequence as 1 to 13 for the top strand (motif 1) according to the basic CRD-binding structure (Fig. 2I) and used subscribed T_1_, T_2_ and T_3_ for the triple-T element of the bottom strand (motif 2) (Fig. 3B).

The axes of the two stems, the 6-bp stem S1 and the 4-bp stem S2, are in a L-shaped ∼90° sharp turn (Fig. 3B). The base pairs in the two stem regions obey the Watson-Crick hydrogen bonding patterns (Fig. 3, D, E and N-Q). The sharp turn of the helix is mediated by the 5-nt bulged loop. Interestingly, the bulge contains an intra-strand C3:G6 base pair (Fig. 3F), which perfectly stacks with the last base pair of stem S1 (Fig. 3G). For the two nucleotides between C3 and G6, C4 protrudes from the bulge (Fig. 3H) and T5 stacks on the other side of the C3:G6 base pair (Fig. 3I). DNA binding assays revealed that the C3:G6 base pair cannot be changed to the three other base pairs (Fig. S5, ss-138 to ss-140; Fig. S8D), but the two nucleotides (C4 and T5) between C3 and G6 can be substituted with any nucleotides (Fig. S6, ss-58 to ss-63).

The last nucleotide of the bulge, T7, is surrounded by two thymine residues of the triple-T element of the opposite strand: stacking with T_1_ (Fig. 3J) and making a single hydrogen bonding (H-bond) with T_2_ (Fig. 3K). The T7 and T_2_ mismatched bases stack with the first base pair of stem S2 (Fig. 3L). The last thymine residue, T_3_ of the triple-T element, locates in the minor groove side of stem S2 and spans the first two base pairs of the CGCT tetranucleotide sequence (Fig. 3M).

After the 4-bp stem S2 (Fig. 3, N-Q), the first purine mismatch at position 12 forms a non-canonical base pair via two hydrogen bonds (Fig. 3R). Like A:T base pair possessing two H-bonds, the G:A mismatch has the same thermal stability (46). The second purine mismatch has their separate ways, with one (A13) staying stacked with the neighboring bases and the other (G13) flipping out and stacking with the side chain of Trp301 (Fig. 3S). Subsequently, the inter-strand sugar-sugar distance decreased to 3.8 Å from that of ∼10.5 Å in the stem B-DNA. In addition, divalent metal ions (Mg^2+^ used in the crystallization) bind between the phosphate groups of close apposition of DNA strands as well as bridge between two unpaired bases (Fig. S10, A-C). The *C2’* atoms of deoxyribose rings, particularly those in the non-base paired regions (B1 and B2), are in van der Waals contacts with other bases or protein side chains (Fig. S10, D-F), suggesting that relief from the steric repulsion of the ribose 2’-OH group in RNA can allow non-B DNA to fold on its own and/or interact with CDCA7.

### Three-zinc-containing DNA-binding domain

Unlike any other DNA-binding domains, the CDCA7 CRD adopts a ‘cross-braced’ topology of three Zn^2+^-coordinating residues by 11 cysteines and one histidine (Fig. 4, A and B; the Zn^2+^-coordinating residues are highlighted and labeled in Fig. 1A). The first zinc ion (Zn1) is coordinated by two CXXC motifs, C_281_HQC_284_ and C_308_GPC_311_. His282, immediately following Cys281, points to the opposite direction and forms a second set of zinc coordination (Zn2) with C_338_XC_340_XXC_343_. The third zinc ion (Zn3) is organized by C_295_XXXXC_300_ and C_331_XXC_334_. The three sets of zinc coordination residues are interconnected: His282 connects Zn1 and Zn2, the three-residue linker between Cys334 and Cys338 links Zn3 to Zn2, and Gly305 is part of the linker between Cys300 (for Zn3) and Cys308 (for Zn1). Gly305, an ICF3 mutated residue, sits right next to Zn3, with an interatomic distance of 4.5 Å (Fig. 4C). The substitution of Gly305 with valine (G305V) would result in repulsion and disruption of Zn3 binding. The cross-braced three-zinc coordination folds the CDCA7 CRD into a globular domain with five short helices and two short strands (Fig. 4D). The unique zinc coordination by CDCA7 differs from the three-zinc-ion-coordinating histone-binding ADD domain in ATRX and DNMT3 family members (Fig. S11).

**Fig. 4.**
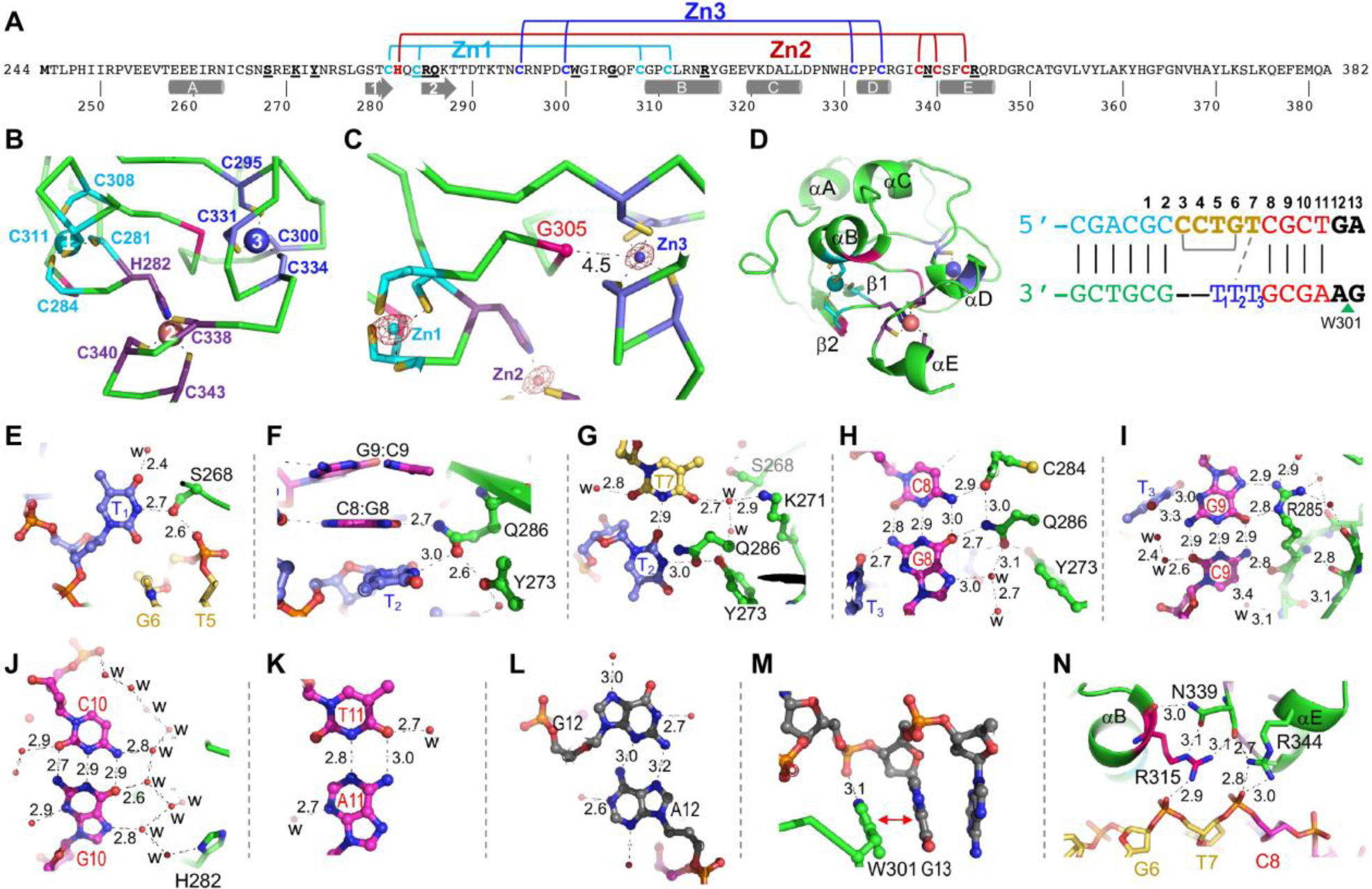
mCDCA7-DNA interactions. (**A**) Cross-brace topology of three Zn binding. Bold Residues involved in chelating Zn and DNA binding are underlined. (**B**) Three zinc atoms are coordinated by 11 cysteines and one histidine. (**C**) Gly305, an ICF3 mutated residue, sits right next to Zn3 and links Zn1 and Zn3. (**D**) A globular CRD with five short helices and two short strands. The residues after helix E are disordered. For convenience, the DNA sequence and its numbering are shown again. (**E**) Ser268 interacts with T_1_. (**F**) Gln286 bridges with T_2_ and G8. (**G**) Gln286 interacts with T_3_. (**H**) The main chain carbonyl oxygen of Cys284 and Gln286 interact with C:G base pair at DNA position 8. (**I**) Arg285 interacts with G:C base pair at DNA position 9 via both side chain and main chain atoms. T_3_ provides two additional H-bonds with G9 at the DNA minor groove. (**J**) Water-mediated interactions with C:G base pair at position 10. (**K**) water-mediated interactions with T:A base pair at position 11. (**L**) A purine mismatch at DNA position 12. (**M**) Trp301 stacks with G12 and forms a H-bond with the phosphate group. (**N**) Arg315 and Arg344 interact with two neighboring DNA phosphate groups.

### DNA base-specific recognition

The triple-T element and the tetranucleotide stem S2 provide most of the functionally important interactions with CDCA7. We made the following observations. First, the *N3* atom of T_1_ along the Watson-Crick edge makes a H-bond with Ser268, which in turn interacts with the phosphate group between T5 and G6 of the opposite strand (Fig. 4E). Second, Gln286 spans two neighboring stacking bases, T_2_ and the Gua of the C:G base pair at position 8 (Fig. 4F). Like T_1_, the Gln286-mediated interaction with T_2_ is via the *N3* atom of the Watson-Crick edge (Fig. 4G). Third, the CpG dinucleotides of stem S2 have the most extensive interactions in both the major and minor grooves. On the major groove side, the C:G base pair at position 8 has Gln286 interaction with the *O6* atom of Gua and the main chain carbonyl oxygen atom of Cys284 interaction with the *N4* atom of Cyt (Fig. 4H). We note that the side chain of Gln286 has saturated potential of its ability to form H-bonds: its amide nitrogen atom and carbonyl oxygen atom each having two H-bonds. On the minor groove side, T_3_ of the triple-T element provides a H-bond with the *N2* atom of Gua. Fourth, the G:C base pair at position 9 has Arg285 on the major groove side, forming the bident H-bonds with the Gua (via the guanidinium group) and an H-bond with the *O4* atom of Cyt (via the main chain carbonyl oxygen) (Fig. 4I). The Arg-Gua interaction is common in specific Gua recognition, but it is unique to have an Arg residue (involving both side-chain and main-chain atoms) interaction with both bases of a G:C base pair (Fig. S10, G-I). The ICF3 mutation of Arg285-to-His or -Cys (R285H/C) would certainly disrupt the Gua recognition. On the minor groove side, T_3_ provides two additional H-bonds with the *N2* and *N3* atoms of Gua. Emphasizing these interactions is the fact that the next two base pairs of stem S2, C:G at position 10 and T:A at position 11, as well as the purine mismatch at position 12, do not conduct direct interactions with CDCA7, instead forming extensive water-mediated interactions (particularly C:G at position 10) (Fig. 4, J-L). Finally, Trp310 forms an aromatic stack interaction with the extra helical Gua at position 13 as well as a H-bond with the associated phosphate group (Fig. 4M). As shown by the DNA-binding assays, an intra-strand C3:G6 base pair of the bulged loop cannot be altered by the three other base pairs. We observed a water-mediated interaction between G6 and Arg269, which also stacks with T_1_ (Fig. S10D).

In addition to the base-specific interactions, CDCA7 contacts ten phosphate groups, including Arg315 interaction with the phosphate group between G6 and T7 and Arg344 interaction with the phosphate group between T7 and C8 (Fig. 4N). The two long side-chain conformations of arginine residues are further stabilized by Asn339 (Fig. 4N), enhancing the Arg-DNA phosphate contacts. The ICF3 mutation of R315H, the substitution of Arg315 by a shorter His side chain, would be disruptive for DNA binding. In summary, CDCA7 residues associated with Zn1 binding, Cys284-Arg285-Qln286, Arg315 between Zn1 and ZF2 binding, and Zn3-associated Arg344 provide the most functionally important interactions in recognizing the DNA bases of the CpG dinucleotides as well as DNA phosphate backbone.

### One of the two motifs that form the non-B DNA is highly enriched at the centromeres of human chromosomes

The basic non-B DNA structure bound by the CDCA7 CRD is formed by two sequence motifs: a 13-mer motif 1 (5’-XYCNNGTCGXYNN-3’) and an 11-mer motif 2 (5’-NNXYCGTTTXY-3’) (Fig. 2I). The two motifs that come together to form the structure do not necessarily need to be continuous or adjacent to each other. We searched the two motifs in the complete human genome assembly T2T-CHM13v2.0 (47,48), including the recently assembled Y chromosome (49). Motif 1 was distributed throughout the genome, with modest enrichment toward one or both ends of some chromosomes. Motif 2, strikingly, showed a prominent peak on each chromosome at the centromere, as evidenced by CENP-A occupation (Fig. 5, Fig. S12). Human centromeres are defined by αSat, an AT-rich repeat family composed of ∼171-bp monomers, which can occur either as large arrays of higher-order repeats (HORs) that are composed of different monomer variants or stretches of divergent monomers (48,50). Annotations of satellite repeats (48,49) confirmed that the centromeric peaks were present in αSat, mostly in active HORs (Fig. 5, Fig. S12), where kinetochore proteins bind (51,52). Indeed, the CGTTT sequence of motif 2, which provides most of the functionally important interactions with CDCA7 (Fig. 4), is present in many αSat sequences deposited in GenBank. Motif 1 was depleted at motif 2 peaks at centromeres (Fig. 5, Fig. S12), perhaps due to the highly biased sequence of the αSat.

**Fig. 5.**
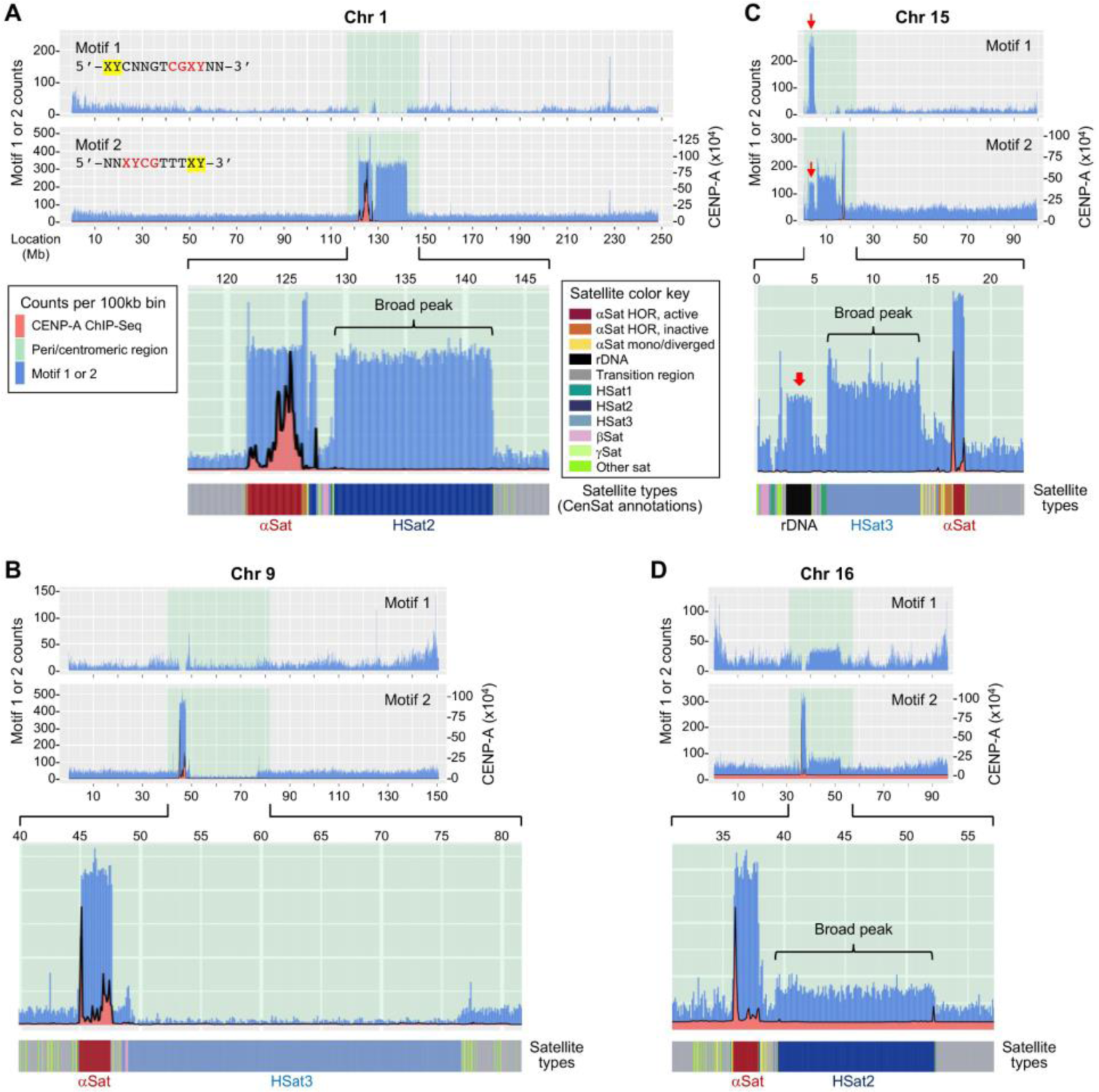
Motif 2 of the non-B DNA is highly enriched at centromeres of human chromosomes. Motifs 1 and 2 that form the non-B DNA (see Figure 2I) were searched in the T2T_CHM13v2.0 human genome assembly. Shown are the results of chromosome 1 (**A**), chromosome 9 (**B**), chromosome 15 (**C**), and chromosome 16 (**D**). The results of other chromosomes are shown in Fig. S12. Top panels: chromosome-wide views of motifs 1 and 2 counts (blue). Highlighted in light green are the peri/centromeric regions. X axis, chromosome locations (Mb); Y axis on the left, motif 1 or 2 counts; Y axis on the right, CENP-A ChIP signal (orange, x10^4^). Middle panel: 5x expansion of the peri/centromeric regions of the motif 2 panel (the Y axis is not proportionally magnified). Bottom panel: CenSat annotations of satellite types in peri/centromeric regions.

In addition to dramatic centromeric enrichment, broad peaks of motif 2 were observed in the pericentromeric regions of chr 1, 15 and, less abundantly, chr 16 (Fig. 5, A, C and D). They were present in HSat2 (chr 1 and 16) and HSat3 (chr 15), respectively (Fig. 5, A, C and D). The broad motif 2 peaks on chr 1 and 15, like centromeric peaks, were accompanied by motif 1 depletion (Fig. 5, A and C), whereas motifs 1 and 2 were both slightly enriched in the pericentromeric region of chr 16 (Fig. 5D). A notable exception is HSat3 in the pericentromeric region of chr 9, which is frequently hypomethylated in ICF syndrome (22), showed depletion of motif 2 (Fig. 5B).

The five acrocentric chromosomes (chr 13, 14, 15, 21 and 22) formed a unique group in that each had a major peak of both motifs 1 and 2 – with motif 1 being more abundant – at the same location in the pericentromeric region, which coincides with rDNA repeats (Fig. 5C, Fig. S12, K, L, Q and R, indicated by red arrows). Chr 14 and 22 each had an additional peak of motifs 1 and 2 at the very end of the p arm, present in satellite DNA annotated as ‘other satellite’ (Fig. S12, L and R, indicated by blue arrows). Analysis of the co-localized motifs 1 and 2 peaks on the acrocentric chromosomes revealed considerable numbers of paired motifs (Table S2), suggesting abundant opportunities for neighboring motifs 1 and 2 to form the non-B DNA structure in these regions.

On chr Y, both motifs 1 and 2 showed similar distribution patterns – one major peak at the very end of the p arm (indicated by blue arrows in Fig. S12T), two major peaks in the pericentromeric region (indicated by red arrows), and many peaks in the large heterochromatic q12 region – with motif 2 being consistently more abundant than motif 1. Satellite annotations revealed that they were present in gamma satellite (γSat) (p-arm end), ‘other satellite’ (pericentromeric region), and HSat3 (q12 region), respectively (Fig. S12T).

Examination of the 4-bp stem S2 in peri/centromeric regions revealed chromosome-specific sequence preferences. For example, chr 14 and 16 strongly prefer CGCT and CGAT, respectively, in motif 1 (Table S3), and most chromosomes prefer CCCG, TCCG and TTCG in motif 2 (Table S4). We note that the CCCGTTT sequence of motif 2 is identical to part of the 17-bp CENP-B box (CTTCGTTGG<u>AAACGGG</u>A) in αSat (53).

### Strand-specific CpG methylation in the non-B DNA has opposite effects on CDCA7 binding

Stem S2 of the non-B DNA contains a CpG dyad that is essential for CDCA7 binding (Fig. 2I, Fig. 4, H and I). Thus, we assessed the effects of CpG methylation. By annealing two single-stranded oligos with or without methylated cytosine (depicted as M), we generated probes that were unmethylated, fully methylated (on both strands), or hemi-methylated on either the forward strand (motif 1-containing Hemi-F) or reverse strand (motif 2-containing Hemi-R) (Fig. 6A). As revealed by EMSA, the mCDCA7 CRD failed to bind the fully methylated and Hemi-R probes but showed stronger binding to the Hemi-F probe, compared to the unmethylated probe (Fig. 6A).

**Fig. 6.**
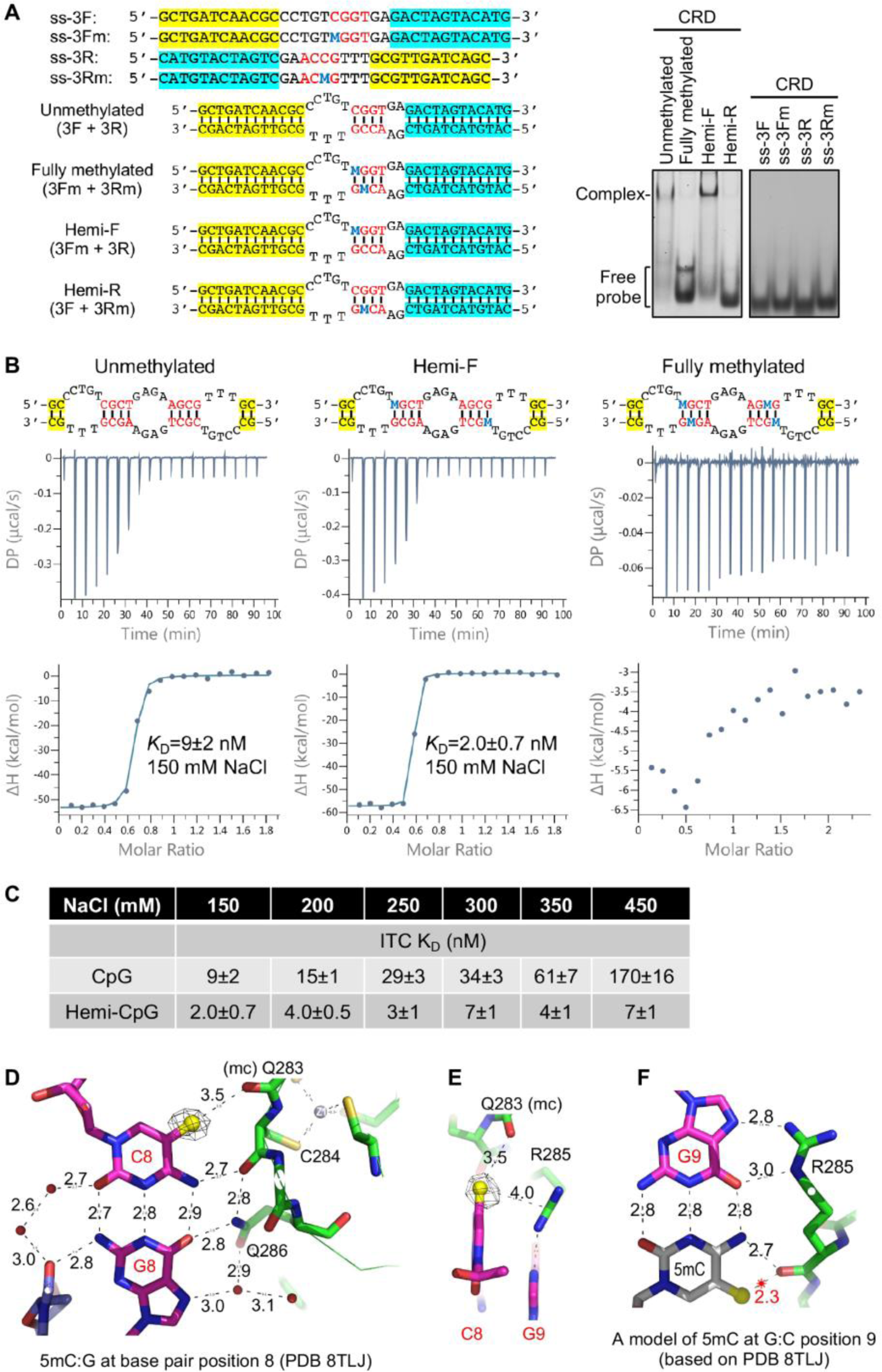
Strand-specific CpG methylation of stem S2 shows the opposite effects on CDCA7 binding. (**A**) EMSA data showing that mCDCA7 CRD-mediated binding of the non-B DNA is enhanced by methylation of the forward strand of stem S 2 (hemi-F probe) but abolished by methylation of the reverse strand (hemi-R probe) or both strands (fully methylated probe). Methylated cytosine (M) is shown in blue. Note that the ss oligos, before annealing, failed to be bound (right). (**B**) ITC assays with similar probes, which confirm that mCDCA7 CRD binds the hemi-F probe with higher affinity (*K*_D_ = 2.0 ± 0.7 nM) compared to that of the unmethylated probe (*K*_D_ = 9 ± 2 nM) but does not bind the fully methylated probe. (**C**) ITC assays with increased salt concentrations, which show that the binding affinity of the unmethylated probe decreases with increasing concentrations of NaCl, whereas the binding affinity of the hemi-F probe is not affected by as high as 450 mM NaCl. (**D**) Structure of mCDCA7 in complex with methylated cytosine at position 8. The methyl group is recognized by the main chain carbonyl oxygen of Gln283. (**E**) A methyl-Arg-Gua triad involving the neighboring 5mC and Gua of the same DNA strand. (**F**) A model of cytosine methylation at position 9 resulting in repulsion with Arg285.

To confirm the effects of methylation, we performed ITC using similar probes. Indeed, the binding affinity of Hemi-F (∼2 nM) was higher than that of the unmethylated probe (∼9 nM) at the condition of 150 mM NaCl, whereas the fully methylated probe failed to be bound by the mCDCA7 CRD (Fig. 6B). To further verify the results, we measured the binding affinities under increased ionic strengths (150-450 mM NaCl). While the binding affinities of the unmethylated probe decreased with increases in ionic strengths (*K*_D_ went from ∼9 nM at 150 mM NaCl to ∼170 nM at 450 mM NaCl), the binding affinities of the Hemi-F probe remained unchanged (*K*_D_ ∼2-7 nM under the concentrations tested) (Fig. 6C, Fig. S7E).

Next, we determined the structures of the CRD of mCDCA7 and hCDCA7, respectively, in complex with hemi-methylated DNA in the context of 32 nt (Table S1). Structural data indicated that the methyl group of 5-methylcytosine (5mC) at base pair position 8 (Hemi-F) is accommodated by forming a van der Waals contact and a weak C-H•••O type H-bond (3.5 Å) with the main-chain carbonyl oxygen of Gln283 (Fig. 6D). In addition, the methyl group of 5mC makes a van der Waals contact with Arg285 which interacts with the neighboring Gua of the same DNA strand (Fig. 6E). In addition, we modeled 5mC onto the cytosine of the opposite strand at the next G:C base pair (Fig. 6F). The methyl group is placed as close as 2.3 Å to the main-chain carbonyl oxygen of Arg285, which is likely to cause steric obstruction with Arg285 in this conformation, perhaps explaining the lack of binding to cytosine methylation at base pair position 9 (Hemi-R).

### Hemi-F, but not Hemi-R, inhibits CDCA7 foci formation and induces hypomethylation of centromeric satellite DNA

Collectively, our findings suggest that the CDCA7 CRD would preferentially bind a specific non-B DNA with stem S2 being hemi-methylated on the forward strand (motif 1), which perhaps contributes to CDCA7 enrichment in constitutive heterochromatin. We asked whether exogenous Hemi-F would affect CDCA7 foci formation. Different amounts of the Hemi-F, Hemi-R or unmethylated probe (Fig. 6A) were transfected into NIH3T3 cells stably expressing HA-mCDCA7 (Fig. 1C). IF analysis at 48h post-transfection revealed that Hemi-F, but not Hemi-R or unmethylated probe, inhibited the enrichment of mCDCA7 in heterochromatin foci (Pattern A) in a dose-dependent manner (Fig. 7, A and B).

**Fig. 7.**
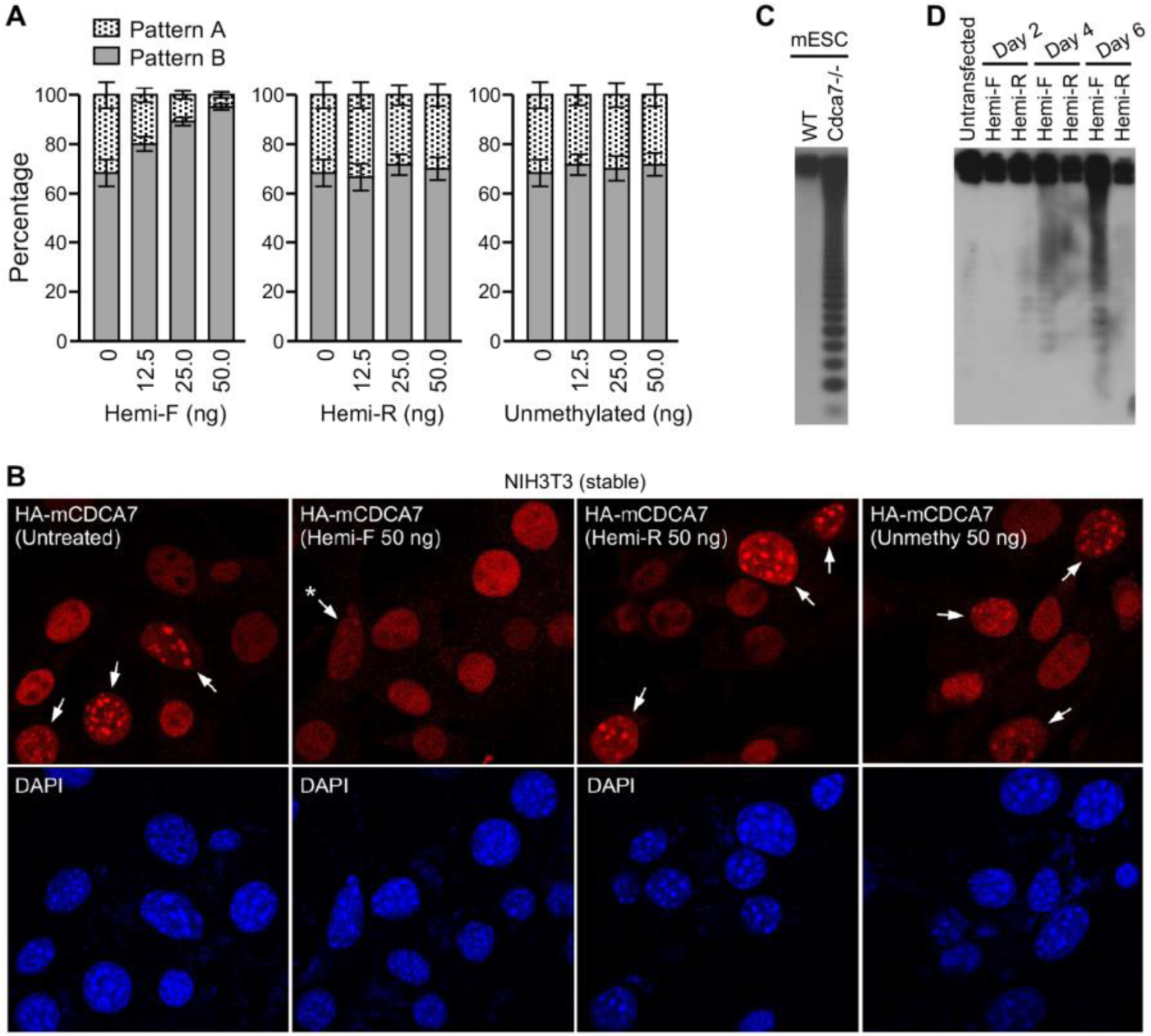
Exogenous Hemi-F inhibits CDCA7 foci formation and induces hypomethylation of minor satellite repeats in murine cells. (**A** and **B**) IF data showing that transfection of the Hemi-F, but not Hemi-R or unmethylated, probe in NIH3T3 cells stably expressing HA-mCDCA7 results in reduced numbers of cells with CDCA7 heterochromatin foci (Pattern A). IF was performed 48 h after transfection of the probes. Shown are the percentages of the two localization patterns (mean ± SD from three experiments) (**A**) and representative images (**B**). (**C**) Southern blotting analysis of genomic DNA from WT and *Cdca7^-/-^* mESCs after digestion with the methylation-sensitive restriction enzyme *Hpa*II, which shows substantial hypomethylation of minor satellite repeats in *Cdca7*-deficient cells (as evidenced by the smear on the gel, which indicates digestion of minor satellite DNA into smaller fragments). (**D**) Southern blot showing that repeated transfections of the Hemi-F, but not Hemi-R, probe in WT mESCs induce loss of methylation at minor satellite repeats.

We previously showed that disruption of *Cdca7* in mESCs results in substantial hypomethylation of the minor satellite repeats in centromeric regions (37) (Fig. 7C). Thus, we assessed the effects of Hemi-F and Hemi-R on DNA methylation by repeatedly transfecting the probes (once every other day) into WT mESCs. As shown in Fig. 7D, Hemi-F, but not Hemi-R, induced detectable loss of methylation at the minor satellite repeats after two times of transfection (day 4) and more obvious hypomethylation after three times of transfection (day 6). These results support the idea that CRD-mediated recognition of hemi-methylated non-B DNA formed at centromeres contributes to the specificity of CDCA7 in regulating DNA methylation.

## Discussion

Repetitive sequences, which are prone to giving rise to non-B DNA (54), are major components of eukaryotic genomes, making up, for instance, over 50% of the human genome (47–49). Thus, the potential of genomic DNA to form non-B structures is enormous. Despite decades of investigations, the biological effects of non-B-form DNA, as well as the molecular mechanisms involved, are not well characterized. In this study, we demonstrate that the CDCA7 CRD forms a novel DNA-binding domain that recognizes a CpG-containing non-B DNA, providing a plausible explanation for the functional specificity of CDCA7 in the regulation of DNA methylation and shedding light on the pathogenesis of ICF syndrome. Unlike two other known methyl-CpG binding domains – the SET and RING-associated (SRA) domain of UHRF1, which binds specifically hemi-methylated CpG (55), and the methyl-CpG-binding domain (MBD) of MeCP2, which binds fully methylated CpG (56) – the CRD of CDCA7 preferentially binds strand-specific hemi-methylated CpG in the context of a non-B DNA. Furthermore, the SRA domain uses base flipping for binding an extra-helical 5mC in an aromatic cage, whereas the MBD and the CDCA7 CRD bind the intra-helical 5mC using an arginine residue in a methyl-Arg-Gua triad (Fig. 6E).

Hypomethylation of satellite repeats, most notably HSat2 on chr 1 and 16, HSat3 on chr 9, and αSat in centromeric regions (22), is considered the primary defect in ICF syndrome, which presumably underlies other cellular and clinical features, such as centromeric instability and antibody deficiency. Thus, elucidating the roles of ICF-related genes – *DNMT3B, ZBTB24, CDCA7* and *HELLS* – in the regulation of DNA methylation is fundamentally important for understanding the pathophysiology of the disease. Previous studies suggest that CDCA7, by recruiting HELLS to heterochromatin, is a key player that controls DNA methylation at satellite repeats (33,36,37). However, little is known about how CDCA7 specifically recruits HELLS to heterochromatin. Based on our findings, we propose that CRD-mediated binding of the CpG-containing non-B DNA confers CDCA7 the specificity to regulate DNA methylation. First, the CDCA7 CRD binds the non-B DNA in a highly specific manner, and the identified ICF3 missense mutations – all in the CRD – disrupt the binding. Second, the peri/centromeric regions in human cells have great potential to form the specific non-B DNA. Bioinformatic analysis of the two sequence motifs in the complete human genome assembly (47–49) revealed that, while motif 1 is distributed (more or less evenly) throughout the genome, motif 2 is highly enriched in centromeric αSat of all chromosomes and is also abundant in HSat2 in the pericentromeric regions of chr 1 and 16, two of the chromosomes most frequently affected in ICF syndrome (Fig. 5, Fig. S12). Intriguingly, some motif 2s are present in the 17-bp CENP-B box (53). *In vitro* evidence suggests that CpG methylation in the CENP-B box inhibits CENP-B binding (57). It would be interesting to determine the possible interplay between CDCA7 and CENP-B in αSat methylation, as well as centromere formation and functions. Third, WT CDCA7, but not ICF3 mutants, is concentrated in constitutive heterochromatin foci during DNA replication, when negative supercoiling and ssDNA are created, both favoring the formation of non-B DNA structures (54). Given that the sequences of satellite DNA repeats are not conserved in the mouse and human genomes and, yet, CDCA7 foci were detected in both mouse and human cells (Fig. 1, Fig. S2), we speculate that the sequence motifs that form the non-B DNA are also abundant in heterochromatin regions in the mouse genome. As an example, we found that a 23-nt sequence on mouse chr 3 could be bound by mCDCA7 CRD *in vitro* (Fig. S7F). However, the distribution of the two motifs in peri/centromeric regions in the mouse genome remains to be determined, as many repetitive sequences have yet to be assembled. Finally, when introduced in cells, a hemi-methylated non-B DNA (Hemi-F) that is preferentially bound by the CRD can inhibit the formation of CDCA7 foci and induce hypomethylation of centromeric satellite repeats, whereas a similar non-B DNA (Hemi-R) that cannot be bound by CDCA7 shows no effects (Fig. 7). The results support the notion that CRD-mediated binding of the specific hemi-methylated non-B DNA contributes to the concentration of CDCA7 in constitutive heterochromatin during DNA replication.

Our observation that CDCA7 CRD-mediated binding of the non-B DNA is differentially affected by strand-specific asymmetric CpG methylation (Fig. 6) provides insights into the mechanism involved in methylation of peri/centromeric satellite repeats. Based on our results, CDCA7 would preferentially bind the non-B DNA structure formed with methylated motif 1 and unmethylated motif 2. One scenario is that, when motif 1, from neighboring and/or distant genomic regions, and motif 2 form the non-B DNA structure, the motif-1 CpG methylation marks would serve as templates for the methylation of motif 2 in peri/centromeric regions. We envisage two related roles for CDCA7: i) temporarily stabilizing the non-B structure; and ii) recruiting HELLS to peri/centromeric regions, where it performs chromatin remodeling and/or recruits components of the DNA methylation machinery to facilitate DNA methylation (28–31). As fully methylated non-B DNA would disrupt the binding by CDCA7, the 5mC ‘templates’ in motif 1 and the methylation machinery could be propagated to neighboring motif 2s. Conceivably, the methylation marks deposited in motif 2s may also spread to neighboring CpG sites, further contributing to the efficiency in methylating satellite repeats. This proliferated process would be severely compromised in ICF syndrome because of DNMT3B inactivation (in ICF1), inhibition of *CDCA7* expression (in ICF2 due to *ZBTB24* mutations), alterations in the CDCA7 CRD (in ICF3), or disruption of *HELLS* (in ICF4). While our results can explain the loss of methylation in most peri/centromeric satellite repeats in ICF syndrome, it is worth noting that motifs 1 and 2 are not enriched in HSat3 on chr 9 (Fig. 5B), which is also hypomethylated in ICF syndrome, albeit to a lesser extent compared to other satellite repeats (22). Given that different types of ICF syndrome show both common and distinct changes in DNA methylation patterns (22,58,59), it is possible that different mechanisms are involved in methylating different regions.

## Material and Methods

### Plasmid constructs

The HA-mCDCA7 construct was described previously (37). The HA-hCDCA7 and HA-mHELLS constructs were generated by cloning synthesized human *CDCA7* cDNA (Accession: NP_665809.1) and PCR-amplified mouse *Hells* cDNA (Accession: NM_008234.3), respectively, into the *pCAG-HA-IRESBlast* vector (60). The GFP-mCDCA7 construct was generated by cloning mouse *Cdca7* cDNA into the *pEGFP-C1* vector (Clontech). The GST-mCDCA7 CRD (pXC2025) and GST-hCDCA7 CRD (pXC2205) constructs were generated by cloning the corresponding CRD fragments (mouse: residues 241-382; human: residues 235-371) into the *pGEX-6P-1* vector (Amersham). Mutations and deletions in mCDCA7 or hCDCA7 were introduced by PCR-based mutagenesis. The primers used and the synthesized human *CDCA7* cDNA are listed in Table S5. All constructs were verified by DNA sequencing.

### Cell culture, transfection, and generation of stable cell lines

NIH3T3, HeLa and HEK293 cells were cultured in Dulbecco’s Modified Eagle Medium (DMEM) supplemented with 10% fetal bovine serum (FBS), 2 mM L-glutamine (Gibco), 50 U/mL penicillin, and 50 μg/mL streptomycin. WT and *Cdca7^-/-^* J1 mESCs (37) were maintained on gelatin-coated petri dishes in serum-containing medium (DMEM supplemented with 15% FBS, 0.1 mM nonessential amino acids, 0.1 mM β-mercaptoethanol, 50 U/mL penicillin, 50 μg/mL streptomycin, and 10^3^ U/mL leukemia inhibitory factor). Transfection was performed using Lipofectamine 2000 (Invitrogen). Stable NIH3T3 cell lines expressing HA-tagged mCDCA7 or the R285H mutant were generated by transfecting the corresponding construct, followed by 7 days of selection with Blasticidin S HCl (Gibco).

### Co-immunoprecipitation (Co-IP) and IF

Co-IP and IF were performed as described previously (61). To map the CDCA7 domain that mediates the interaction with HELLS, HA-mHELLS and GFP-mCDCA7 proteins (full-length, the R315H mutation, and the ΔLZ and ΔCRD deletions) were co-expressed in HEK293 cells, the cell lysates were immunoprecipitated with GFP antibody (Abcam, ab1218), and the precipitated proteins were immunoblotted with HA antibody [Cell Signaling Technology (CST), #3724]. The localization patterns of WT and mutant mCDCA7 and hCDCA7 were determined by IF analysis of NIH3T3 and HeLa cells, respectively, that expressed HA-tagged CDCA7 proteins. Constitutive heterochromatin foci were marked by DAPI-bright spots in NIH3T3 cells or stained with H3K9me3 antibody (CST, #13969) in HeLa cells.

### Cell synchronization and cell cycle analysis

To assess CDCA7 localization patterns during the cell cycle, NIH3T3 cells stably expressing HA-mCDCA7 were synchronized as described previously (62). Briefly, cells were first arrested at the G0/G1 phase by culturing them in DMEM medium with 0.5% FBS for 48 h, then 10% FBS was added to induce the cells to re-enter the cell cycle, and, at different timepoints (6-20 h) following serum induction, some cells were stained with propidium iodide for cell cycle analysis by flow cytometry and other cells were analyzed for CDCA7 localization by IF with HA antibody.

### DNA methylation analysis

Methylation of minor satellite DNA in mESCs was analyzed by Southern blotting after digestion of genomic DNA with the methylation-sensitive enzyme *Hpa*II, as described previously (63,64).

### GST-CDCA7 CRD protein expression and purification

Recombinant GST fusion proteins were expressed and purified as described previously (35). Plasmids expressing mCDCA7 or hCDCA7 CRD fragments were transformed into *Escherichia coli* strain BL21-codon-plus (DE3)-RIL. Bacteria were grown in LB broth at 37℃ until the log phase (OD_600nm_ = 0.4-0.5), when the temperature was lowered to 16℃ and 25 µM ZnCl_2_ was added. When OD_600nm_ reached 0.8, 0.2 mM isopropylthio-β-galactoside (IPTG) was added, followed by continuing growth for 20 h at 16℃. Cells were lysed by sonication in lysis buffer [20 mM Tris-HCl pH 7.5, 250 mM NaCl, 5% glycerol, 0.5 mM tris (2-carboxyethl) phosphine (TCEP), and 25µM ZnCl_2_]. After removal of the debris by centrifugation, the supernatant was loaded onto a 5-mL GSTrap column (GE Healthcare). The resin was washed by lysis buffer and the bound protein was eluted in 100 mM Tris-HCl pH 8.0, 250 mM NaCl, 5% glycerol, 0.5 mM TCEP and 20 mM glutathione (reduced form). For SELEX and EMSA, the GST-CRD fusion proteins were used.

Protein purification was further carried out at 4℃ through a multi-column chromatography protocol for ITC and crystallography. The GST tag was removed by digestion with PreScission protease (produced in-house). The cleaved proteins were dialyzed in a buffer consisting of 20 mM Hepes pH 7.0, 0.1 M NaCl, 0.5% glycerol and 0.5 mM TCEP and then loaded onto columns of HiTrap Q HP (5mL) and HiTrap Heparin HP (5mL) (GE Healthcare) connected in tandem. After washing with the same buffer, the Q column was disconnected, and the target protein was eluted from the Heparin column by 0.1 to 1 M NaCl gradient. For each protein, the peak fractions eluted from the Heparin column were pooled and loaded onto a second GSTrap column, from which the flow-through was collected, concentrated, and loaded onto a HiLoad 16/60 Superdex S200 column (GE Healthcare) equilibrated with 20 mM Tris-HCl pH 7.5, 150 mM NaCl, 5% glycerol and 0.5 mM TCEP. The protein fractions were pooled, concentrated, and stored at −80℃ prior to use.

### SELEX

To identify potential DNA sequences recognized by the CDCA7 CRD, SELEX was performed according to the originally reported procedures (65,66), with modifications. GST-mCRD conjugated on Glutathione Sepharose 4B beads and a synthetic ssDNA library (ThermoFisher Scientific, NC1108024) of random 30-mer sequences flanked by 23 nt at each end (5’-TAG GGA AGA GAA GGA CAT ATG AT-3’ and 5’-TTG ACT AGT ACA TGA CCA CTT GA-3’) were incubated in binding buffer (50 mM Tris-HCl pH 7.4, 150 mM NaCl, 0.1 mg/ml BSA, 3 mM DTT, 20 μM ZnSO4, and 5 μg/ml salmon sperm DNA) for 1 h at room temperature. The beads were washed with binding buffer for three times, and bound ssDNA was eluted in water by boiling for 5 min, followed by snap cooling on ice for 3 min. The DNA was extracted by phenol/chloroform (25 : 24) and used as the template for PCR amplification – with primers complementary to the 23-nt flanking sequences (see Table S5) – to obtain the ssDNA pool for the next round of selection. The PCR products were extracted by phenol/chloroform (25 : 24), heated at 95°C for 5 min and then snap-cooled on ice. To minimize nonspecific binding of DNA species, we applied the counter-selection step using GST-mCRD:R274H mutant before GST-mCRD was used in each cycle. After six rounds of selection, the PCR products were subcloned into the *pCR2.1-TOPO* TA cloning vector (ThermoFisher Scientific, #450641), and 30 clones were sequenced.

### EMSA

GST-CRD fusion proteins (20 nM) were incubated with DNA probes (10 nM) in binding buffer (2.5% glycerol, 1 mM MgCl_2_, 0.5 mM EDTA, 0.5 mM DTT, 0.1 mg/ml BSA, 20 μM ZnSO_4_, 50 mM NaCl, and 10 mM Tris-HCl pH7.5) for 30 min at room temperature. Then the samples were subjected to electrophoresis through 5% native polyacrylamide gel in 0.5x TBE buffer and imaged with 9410 Typhoon variable mode imager (GE Healthcare).

### ITC

The ITC experiments were performed on a Microcal PEAQ-ITC instrument (Malvern) at 25℃. The protein was diluted to 50 μM and dialyzed in a buffer consisting of 20 mM Tris-HCl pH 7.5 and 150 mM NaCl. Then 150 µl protein sample was loaded into syringe. ssDNA was diluted to 5 µM using the same buffer and 300 µl DNA sample was loaded into the sample cell. The titration protocol was the same for all the measurements, which was composed of a single initial injection of 0.2 µl protein, followed by 19 injections of 2 µl protein into DNA samples, the intervals between injections was set to 300s and a reference power is 8 µcal s^-1^. Curve fitting to a single-site binding model was performed by MicoCal PEAQ-ITC.

### Crystallography

The protein-DNA complex was prepared by mixing the purified mCDCA7 CRD (residues 244-382) with the 36-mer, 34-mer, 32-mer, 26-mer or 32-mer 5mC oligonucleotides (annealed in 10 mM Tris-HCl pH 7.5, 50 mM NaCl) in a 1:1.2 ratio following by 1 h incubation on ice. An Art Robbins Gryphon Crystallization Robot was used to set up screens of the sitting drop of 0.4 µl at ∼19℃ via vapor diffusion. The complex crystal with 36-mer oligonucleotides (PDB 8TLE and 8TLF) were obtained under the condition of 0.2 M MgCl_2_, 0.1 M Tris-HCl pH 8.5 and 25% polyethylene glycol (PEG) 3350. The complex crystal with 34-mer (PDB 8TLG) or 32-mer (PDB 8TLH) oligonucleotides were grown under the condition of 0.1 M MgCl_2_, 0.1 M Tris-HCl pH 8.5 and 25% PEG3350. The complex crystal with 26-mer oligonucleotides (PDB 8TLL) were obtained under the condition of 0.1 M MgCl_2_, 0.1 M Tris-HCl pH 8.5 and 23% PEG3350. The complex crystal with 32-mer 5mC oligonucleotides (PDB 8TLJ) were obtained under the condition of 0.1 M MgCl_2_, 0.1 M Bis-Tris pH 6.2 and 25% PGE3350. The complex crystal of hCDCA7 CRD with 32-mer 5mC oligonucleotides (PDB 8TLK) were grown under the condition of 0.2 M MgCl_2_, 0.1 M Bis-Tris pH 6.0 and 25% PEG3350.

Crystals were flash frozen using 20% (v/v) ethylene glycol as the cryoprotectant. Resulting crystallographic datasets were processed with HKL2000 (67). Two scaled files were output with one file combining Bijvoet pairs and the other keeping them separate. The dataset for the first structure (PDB 8TLE) was examined for single-wavelength anomalous dispersion (Zn-SAD) phasing using the PHENIX Xtriage module, which reported severe anisotropy but also a good anomalous signal to ∼4 Å. The PHENIX AutoSol module (68) readily found three zinc atom positions to give an interpretable map with initial figure-of-merit of 0.31 and gave a density-modified map with an R-factor of 0.42 (Table S1). The initial electron density showed recognizable molecular features of the β-strands and α-helices and DNA bases and backbone. Reinserting the zinc positions into AutoSol and utilizing the full resolution of the dataset gave a better map allowing for our initial model build. This resulting structure was utilized for molecular replacement in the PHENIX PHASER module (69) for the other structures. All structure refinements were performed by PHENIX Refine (70), with 5% randomly chosen reflections for validation by R-free values. Manual (re)building with COOT (71) was conducted carefully between refinement cycles. Structure quality was analyzed during PHENIX refinements and validated by the PDB validation server (72). Molecular graphics were generated using open-source PyMOL (http://www.pymol.org/pymol).

### Bioinformatics analysis

The CENP-A ChIP-seq dataset (SRR766736) was downloaded from the GenBank Short Read Archive. T2T centromere/satellite (CenSat) data (chm13v2.0_censat_v2.0.bed) was accessed from the Telomere-to-Telomere (T2T) Consortium CHM13 project site (https://github.com/marbl/CHM13). Motifs 1 and 2 sequences and the CENP-A ChIP sequences were aligned to the T2T genome (T2T-CHM13v2.0), and the aligned counts on both strands were binned at 100 kb to generate the coverage plots showing chromosome-wide motif alignments and CENP-A ChIP fragment alignments. The CenSat annotation map was produced using the CenSat bed file and assigning the color of the annotation to each 25-kb bin region across the peri/centromere regions, except for chr Y where the bins spanned the entire chromosome.

## Acknowledgments

We thank Dr. Richard Wood (MD Anderson Cancer Center) and Dr. Cristina Cardoso (Technische Universität Darmstadt) for discussion, Ms. Yu Cao and Dr. Yang Zeng (MD Anderson Cancer Center) for technical assistance, and Dr. Xiangpeng Kong (New York University) for assistance of access to 17-ID-1 beamtime. We thank the beamline scientists of Southeast Regional Collaborative Access Team (SER-CAT) at the Advanced Photon Source (APS), Argonne National Laboratory and 17-ID-1 of the National Synchrotron Light Source II, Brookhaven National Laboratory.

## Funding

National Institutes of Health grant R01AI1214030 (TC)

National Institutes of Health grant R35GM134744 (XC)

Cancer Prevention and Research Institute of Texas grant RR160029 (XC)

Sam and Freda Davis Fund fellowship (ZY)

National Institutes of Health equipment grants S10_RR25528, S10_RR028976, and S10_OD027000

U.S. Department of Energy contract W-31-109-Eng-38

National Synchrotron Light Source II resources 17-ID-1 under contract DE-SC0012704

## Author contributions

Conceptualization: TC, XC

Investigation: SH, RR, ZY, JRH, BL (Bigang), JD

Bioinformatics analysis: MDB, BL (Bin), YL

Supervision: BL (Bin), XZ, XC, TC

Writing: TC, XC

## Data and materials availability

All data are available in the main text or the supplementary materials. The authors have deposited the X-ray structure (coordinates) and the source data (structure factor file) of CDCA7-DNA to the PDB, and these will be released upon article publication under accession numbers PDB 8TLE (mCDCA7 and oligo 36-nt), PDB 8TLF (mCDCA7 and oligo 36-nt), PDB 8TLG (mCDCA7 and oligo 34-nt), PDB 8TLH (mCDCA7 and oligo 32-nt), PDB 8TLL (mCDCA7 and oligo 26-nt), PDB 8TLJ (mCDCA7 and 5mC oligo in 32-nt) and PDB 8TLK (hCDCA7 and 5mC oligo in 32-nt).

