## Supplemental Figs. S1-S12, Tables S1 and S5 for "The ICF syndrome protein CDCA7 harbors a unique DNA-binding domain that recognizes a CpG dyad in the context of a non-B DNA"

Swanand Hardikar *et al.*

**This PDF file includes:**

Figs. S1 to S12
Tables S1 and S5

**Other Supplemental Information for this manuscript include the following Excel files:**

Tables S2 to S4

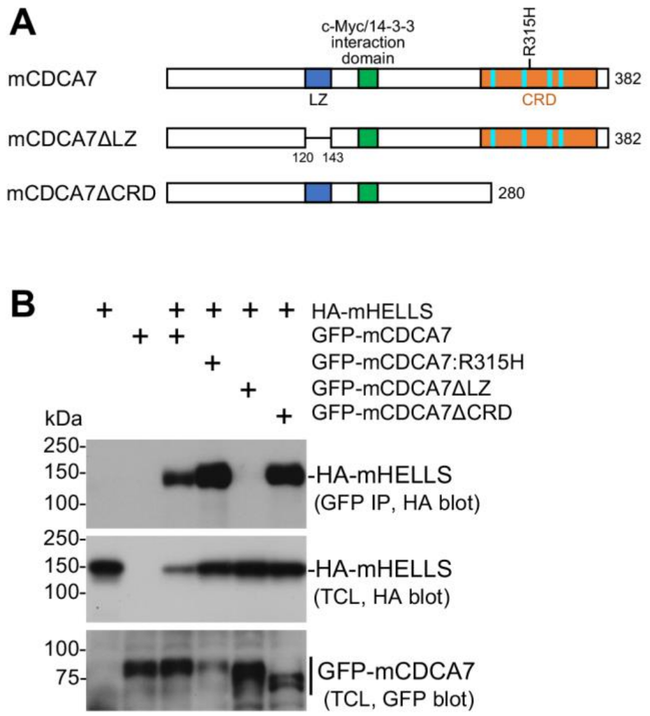


**Fig. S1. The leucine zipper motif of CDCA7 is required for interaction with HELLS.** (**A**) The mCDCA7 constructs used for Co-IP assay. LZ, leucine zipper; CRD, cysteine-rich domain. (**B**) Co-IP results showing that deletion of the leucine zipper motif (ΔLZ) abolishes the interaction between mCDCA7 and mHELLS. HA-tagged mHELLS and GFP-tagged mCDCA7 proteins were co-expressed in HEK293 cells, the total cell lysates (TCL) were immunoprecipitated with GFP antibody, and the precipitated proteins and TCL were immunoblotted with HA or GFP antibody, as indicated.

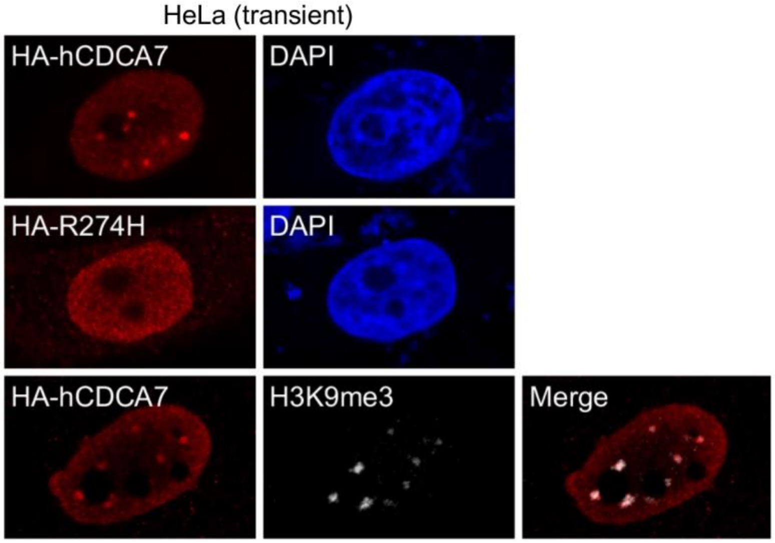


**Fig. S2. hCDCA7, but not the R274H ICF mutant, is enriched in heterochromatin foci in HeLa cells.** IF data showing HA-tagged hCDCA7 and the R274H mutant transiently expressed in HeLa cells. While DAPI-stained heterochromatin foci in human cells are not as prominent as in mouse cells, double IF analysis with HA and H3K9me3 antibodies shows that CDCA7 foci overlap with H3K9me3 foci (bottom), confirming that CDCA7 is enriched in constitutive heterochromatin in human cells.

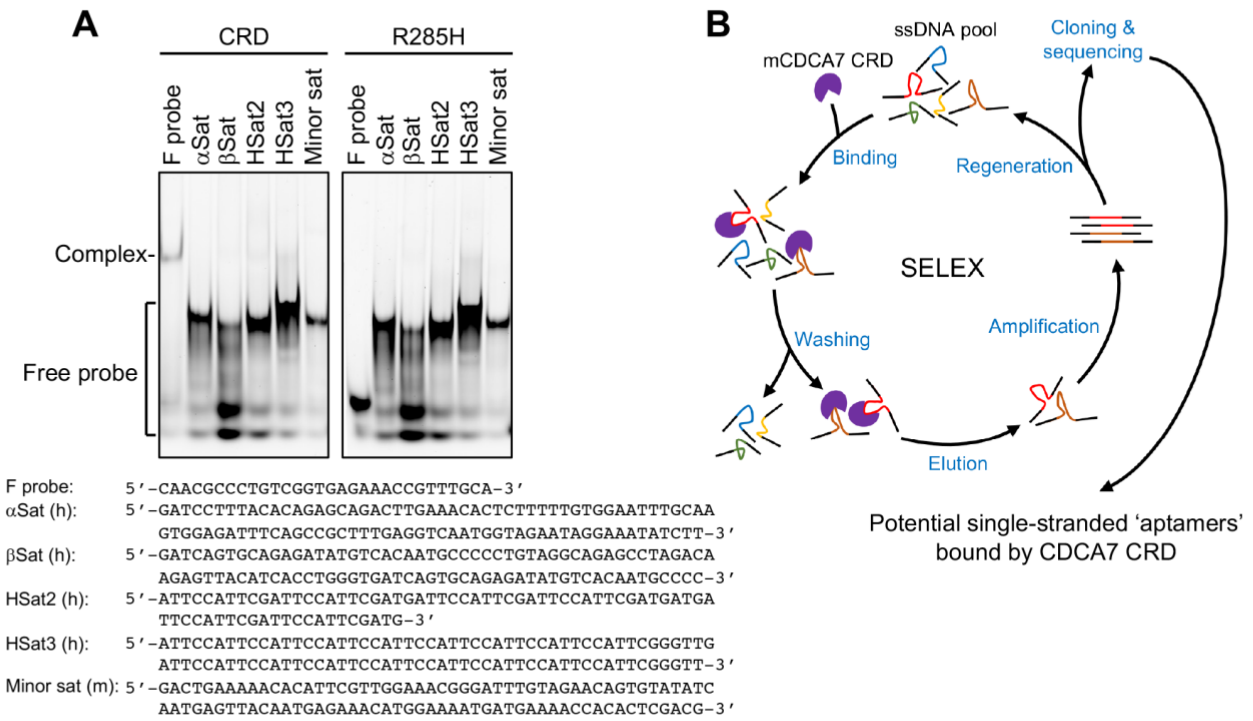


**Fig. S3. Identification of potential DNA sequences bound by CDCA7 using candidate approach and SELEX.** (**A**) EMSA data showing that mCDCA7 CRD fails to bind the repeating units of common satellite DNAs in peri/centromeric regions of human and mouse genomes. The sequences of the single-stranded DNA probes are shown below. As a positive control, the F probe is bound by mCDCA7 CRD, but not by the R285H mutant. (**B**) Schematic representation of the SELEX procedure for identifying DNA sequences specifically or preferentially bound by mCDCA7 CRD.

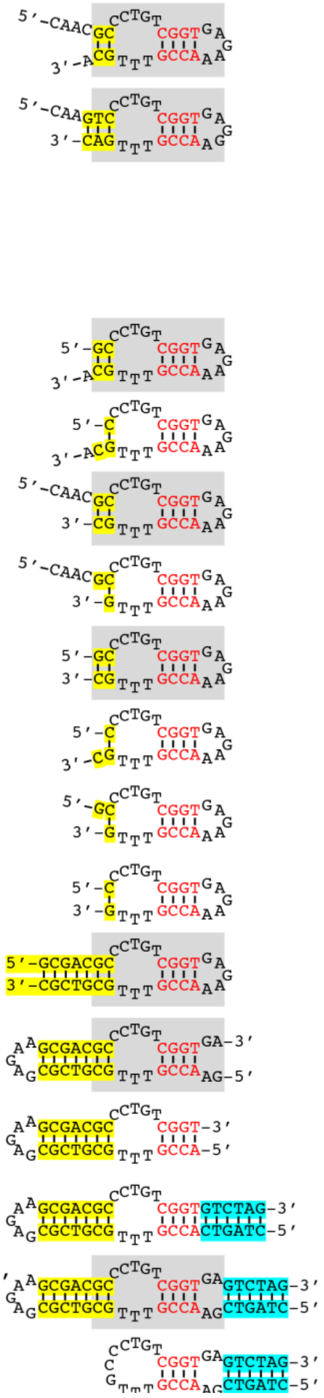


| Probe and sequence (comment) | CRD binding |
|---|---|
| ss-1: 5'-CAACGCCCTGTCGGTGAGAAACCGTTTGCA-3' (Seq-1 identified by SELEX, identical to F probe in Fig. 2) | + |
| ss-2: 5'-CAAGTCCCTGTCGGTGAGGAACCGTTTGAC-3' (Seq-2 identified by SELEX) | + |
| ss-3: 5'-TCGGTGAGAAACCGTTTGCA-3' (10 nt at 5' end of ss-1 deleted) | − |
| ss-4: 5'-CAACGCCCTGACCGTTTGCA-3' (10 nt in the middle of ss-1 deleted) | − |
| ss-5: 5'-CAACGCCCTGTCGGTGAGAA-3' (10 nt at 3' end of ss-1 deleted) | − |
| ss-6: 5'-GCCCTGTCGGTGAGAAACCGTTTGCA-3' (4 nt at 5' end of ss-1 deleted) | + |
| ss-7: 5'-CCCTGTCGGTGAGAAACCGTTTGCA-3' (5 nt at 5' end of ss-1 deleted) | − |
| ss-8: 5'-CAACGCCCTGTCGGTGAGAAACCGTTTGC-3' (1 nt at 3' end of ss-1 deleted) | + |
| ss-9: 5'-CAACGCCCTGTCGGTGAGAAACCGTTTG-3' (2 nt at 3' end of ss-1 deleted) | − |
| ss-10: 5'-GCCCTGTCGGTGAGAAACCGTTTGC-3' (4 nt at 5' end & 1 nt at 3' end of ss-1 deleted, minimal CRD-binding sequence) | + |
| ss-11: 5'-CCCTGTCGGTGAGAAACCGTTTGC-3' (1 nt at 5' end of ss-10 deleted) | − |
| ss-12: 5'-GCCCTGTCGGTGAGAAACCGTTTG-3' (1 nt at 3' end of ss-10 deleted) | − |
| ss-13: 5'-CCCTGTCGGTGAGAAACCGTTTG-3' (1 nt at 5' & 3' ends of ss-10 deleted) | − |
| ss-14: 5'-GCCCTGTCGGTGAGAAACCGTTTGC-3' (CRD-binding sequence with extended stem S1) | + |
| ss-15: 5'-GAACCGTTTGCGTCGCGAGAAGCGACGCCCTGTCGGTGA-3' (reshuffled sequence elements that would form similar non-B DNA) | + |
| ss-16: 5'-ACCGTTTGCGTCGCGAGAAGCGACGCCCTGTCGGT-3' (similar to ss-15, without the two mismatches at 5' and 3' ends) | − |
| ss-17: 5'-CTAGTCACCGTTTGCGTCGCGAGAAGCGACGCCCTGTCGGTGACTAG-3' [similar to ss-16 (see Fig. 3E), with extended stem S2 (without bubble B2)] | − |
| ss-18: 5'-CTAGTCGAACCGTTTGCGTCGCGAGAAGCGACGCCCTGTCGGTGAGACTAG-3' (similar to ss-17, with bubble B2) | + |
| ss-19: 5'-CTAGTCGAACCGTTTGCCCTGTCGGTGAGACTAG-3' (sequence predicted to form similar hairpin as ss-18, without stem S1) | − |

**Fig. S4. Mutagenesis analysis to determine the minimal DNA sequence bound by the CDCA7 CRD.** Binding was determined by EMSA. The grey area is the basic CRD-binding structure (see Fig. 2I).

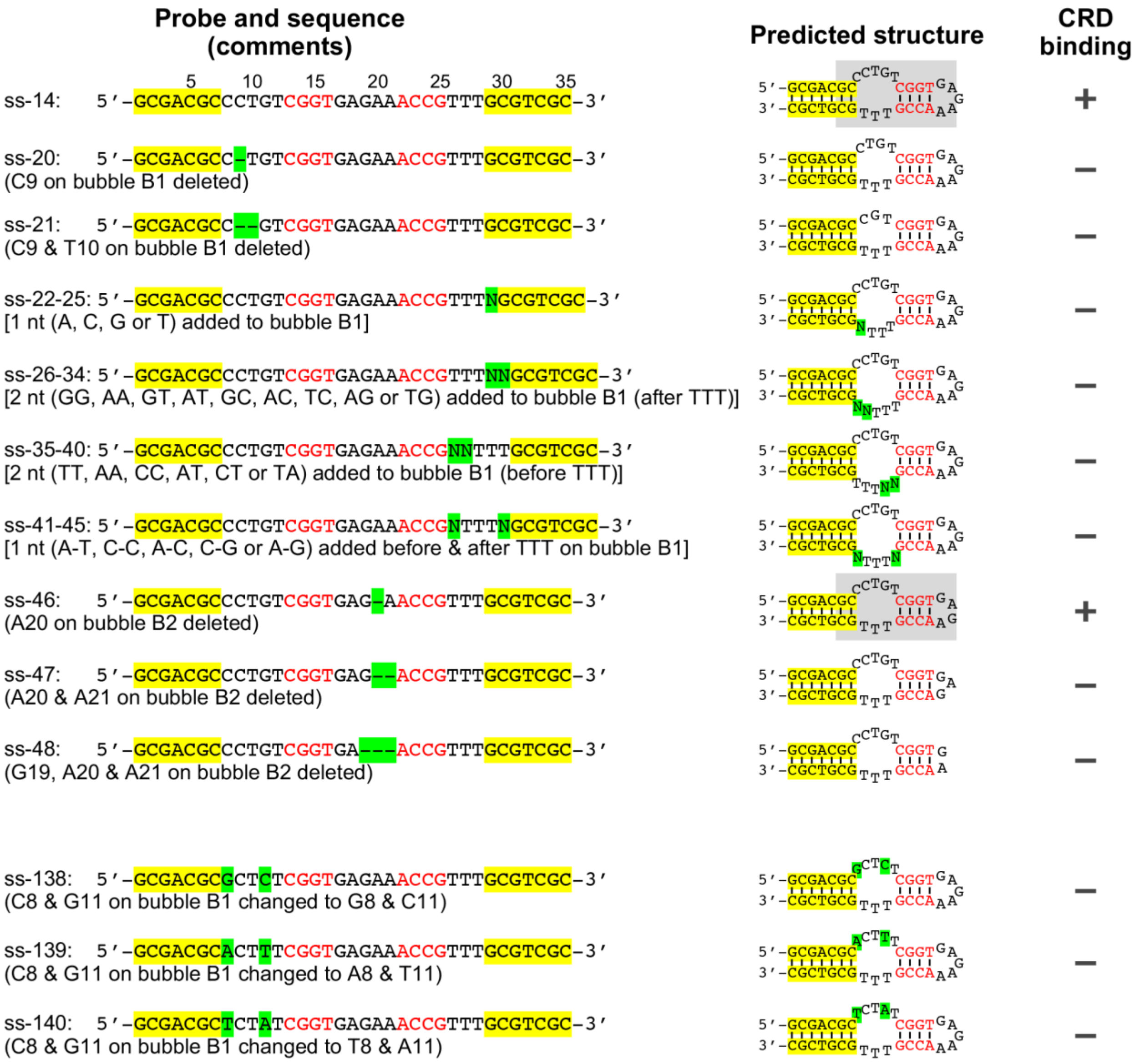


**Fig. S5. Mutagenesis analysis of bubbles B1 and B2 of the non-B DNA structure for binding by the CDCA7 CRD.** Deletions (-), insertions (N) and mutations are highlighted in green. Binding was determined by EMSA. The grey area is the basic CRD-binding structure (see Fig. 2I).

| Probe and sequence (comments) | Predicted structure | CRD binding |
|---|---|---|
| ss-14: 5′-GCGACGCCCTGTCGGTGAGAAACCGTTTGCGTCGC-3′ (positions 5, 10, 15, 20, 25, 30, 35) | 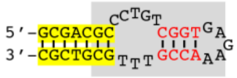 | + |
| ss-49-51: 5′-GCGACNCCCTGTCGGTGAGAAACCGTTTGCGTCGC-3′ (G6 on stem S1 mutated to A, C or T) | 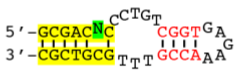 | − |
| ss-52-54: 5′-GCGACGNCCTGTCGGTGAGAAACCGTTTGCGTCGC-3′ (C7 on stem S1 mutated to A, G or T, resulting in larger bubble B1) | 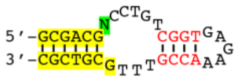 | − |
| ss-55-57: 5′-GCGACGCNCTGTCGGTGAGAAACCGTTTGCGTCGC-3′ (C8 on bubble B1 mutated to A, C or T) | 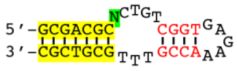 | − |
| ss-58-60: 5′-GCGACGCCNTGTCGGTGAGAAACCGTTTGCGTCGC-3′ (C9 on bubble B1 mutated to A, C or T) | 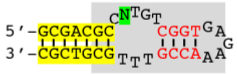 | + |
| ss-61-63: 5′-GCGACGCCCNGTCGGTGAGAAACCGTTTGCGTCGC-3′ (T10 on bubble B1 mutated to A, C or G) | 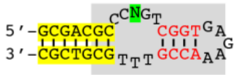 | + |
| ss-64-66: 5′-GCGACGCCCTNTCGGTGAGAAACCGTTTGCGTCGC-3′ (G11 on bubble B1 mutated to A, C or T) | 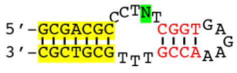 | − |
| ss-67-69: 5′-GCGACGCCCTGNCGGTGAGAAACCGTTTGCGTCGC-3′ (T12 on bubble B1 mutated to A, C or G) | 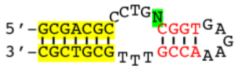 | − |
| ss-70-72: 5′-GCGACGCCCTGTNGGTGAGAAACCGTTTGCGTCGC-3′ (C13 on stem S2 mutated to A, G or T) | 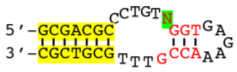 | − |
| ss-73-75: 5′-GCGACGCCCTGTCNGTGAGAAACCGTTTGCGTCGC-3′ (G14 on stem S2 mutated to A, C or T) | 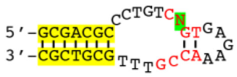 | − |
| ss-76-78: 5′-GCGACGCCCTGTCGNTGAGAAACCGTTTGCGTCGC-3′ (G15 on stem S2 mutated to A, C or T) | 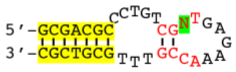 | − |
| ss-79-81: 5′-GCGACGCCCTGTCGGNGAGAAACCGTTTGCGTCGC-3′ (T16 on stem S2 mutated to A, C or G) | 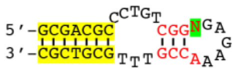 | − |
| ss-82-84: 5′-GCGACGCCCTGTCGGTNAGAAACCGTTTGCGTCGC-3′ (G17 on bubble B2 mutated to A, C or T) | 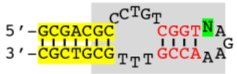 | + |
| ss-85-87: 5′-GCGACGCCCTGTCGGTGNGAAACCGTTTGCGTCGC-3′ (A18 on bubble B2 mutated to C, G or T) | 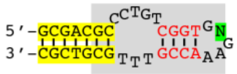 | + |
| ss-88-90: 5′-GCGACGCCCTGTCGGTGANAAACCGTTTGCGTCGC-3′ (G19 on bubble B2 mutated to A, C or T) | 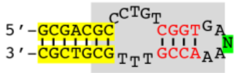 | + |
| ss-91-93: 5′-GCGACGCCCTGTCGGTGAGNAACCGTTTGCGTCGC-3′ (A20 on bubble B2 mutated to C, G or T) | 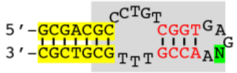 | + |
| ss-94-96: 5′-GCGACGCCCTGTCGGTGAGANACCGTTTGCGTCGC-3′ (A21 on bubble B2 mutated to C, G or T) | 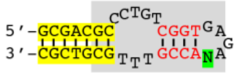 | + |
| ss-97-99: 5′-GCGACGCCCTGTCGGTGAGAANCCGTTTGCGTCGC-3′ (A22 on stem S2 mutated to C, G or T) | 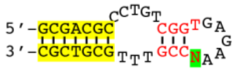 | − |
| ss-100-102: 5′-GCGACGCCCTGTCGGTGAGAAANCGTTTGCGTCGC-3′ (C23 on stem S2 mutated to A, G or T) | 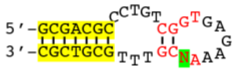 | − |
| ss-103-105: 5′-GCGACGCCCTGTCGGTGAGAAACNGTTTGCGTCGC-3′ (C24 on stem S2 mutated to A, G or T) | 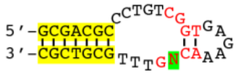 | − |
| ss-106-108: 5′-GCGACGCCCTGTCGGTGAGAAACCNTTTGCGTCGC-3′ (G25 on stem S2 mutated to A, C or T) | 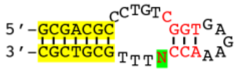 | − |

| Probe and sequence (comments) | CRD binding |
|---|---|
| ss-109-111: 5′-GCGACGCCCTGTCGGTGAGAAACCGNTTGCGTCGC-3′ (T26 on bubble B1 mutated to A, C or G) | − |
| ss-112-114: 5′-GCGACGCCCTGTCGGTGAGAAACCGTNTGCGTCGC-3′ (T27 on bubble B1 mutated to A, C or G) | − |
| ss-115-117: 5′-GCGACGCCCTGTCGGTGAGAAACCGTTNGCGTCGC-3′ (T28 on bubble B1 mutated to A, C or G) | − |
| ss-118-120: 5′-GCGACGCCCTGTCGGTGAGAAACCGTTTNCGTCGC-3′ (G29 on stem S1 mutated to A, C or T, resulting in larger bubble B1) | − |
| ss-121-123: 5′-GCGACGCCCTGTCGGTGAGAAACCGTTTGNGTCGC-3′ (C30 on stem S1 mutated to A, G or T) | − |
| ss-124: 5′-GCGACGACCTGTCGGTGAGAAACCGTTTTCGTCGC-3′ (C7 & G29 on stem S1 changed to A7 & T29) | + |
| ss-125: 5′-GCGACGTCCTGTCGGTGAGAAACCGTTTACGTCGC-3′ (C7 & G29 on stem S1 changed to T7 & A29) | + |
| ss-126: 5′-GCGACGCCCTGTGGGTGAGAAACCCTTTGCGTCGC-3′ (C13 & G25 on stem S2 changed to G13 & C25) | − |
| ss-127: 5′-GCGACGCCCTGTAGGTGAGAAACCTTTTGCGTCGC-3′ (C13 & G25 on stem S2 changed to A13 & T25) | − |
| ss-128: 5′-GCGACGCCCTGTTGGTGAGAAACCATTTGCGTCGC-3′ (C13 & G25 on stem S2 changed to T13 & A25) | − |
| ss-129: 5′-GCGACGCCCTGTCCGTGAGAAACGGTTTGCGTCGC-3′ (G14 & C24 on stem S2 changed to C14 & G24) | − |
| ss-130: 5′-GCGACGCCCTGTCAGTGAGAAACTGTTTGCGTCGC-3′ (G14 & C24 on stem S2 changed to A14 & T24) | − |
| ss-131: 5′-GCGACGCCCTGTCTGTGAGAAACAGTTTGCGTCGC-3′ (G14 & C24 on stem S2 changed to T14 & A24) | − |
| ss-132: 5′-GCGACGCCCTGTCGCTGAGAAAGCGTTTGCGTCGC-3′ (G15 & C23 on stem S2 changed to C15 & G23) | + |
| ss-133: 5′-GCGACGCCCTGTCGATGAGAAATCGTTTGCGTCGC-3′ (G15 & C23 on stem S2 changed to A15 & T23) | + |
| ss-134: 5′-GCGACGCCCTGTCGTTGAGAAAACGTTTGCGTCGC-3′ (G15 & C23 on stem S2 changed to T15 & A23) | + |
| ss-135: 5′-GCGACGCCCTGTCGGGGAGAACCCGTTTGCGTCGC-3′ (T16 & A22 on stem S2 changed to G16 & C22) | + |
| ss-136: 5′-GCGACGCCCTGTCGGCGAGAAGCCGTTTGCGTCGC-3′ (T16 & A22 on stem S2 changed to C16 & G22) | + |
| ss-137: 5′-GCGACGCCCTGTCGGAGAGAATCCGTTTGCGTCGC-3′ (T16 & A22 on stem S2 changed to A16 & T22) | + |

Predicted structure

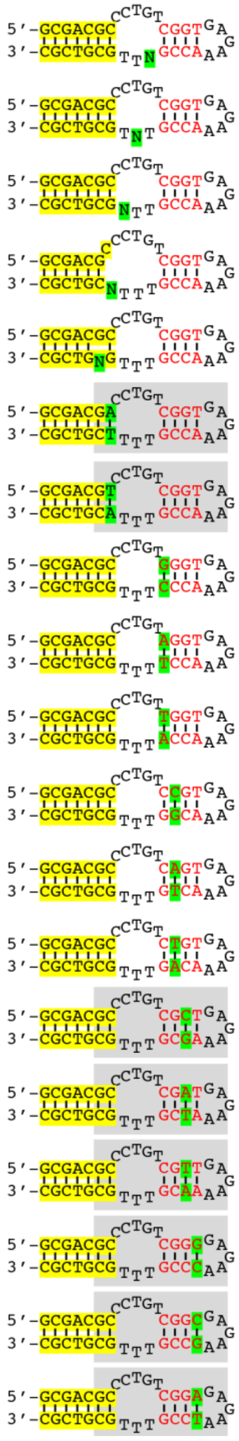

**Fig. S6. Mutagenesis analysis to determine the nucleotides in the non-B DNA structure that are essential for CDCA7 CRD binding.** Mutations are highlighted in green. Binding was determined by EMSA. The grey area is the basic CRD-binding structure (see Fig. 2I).

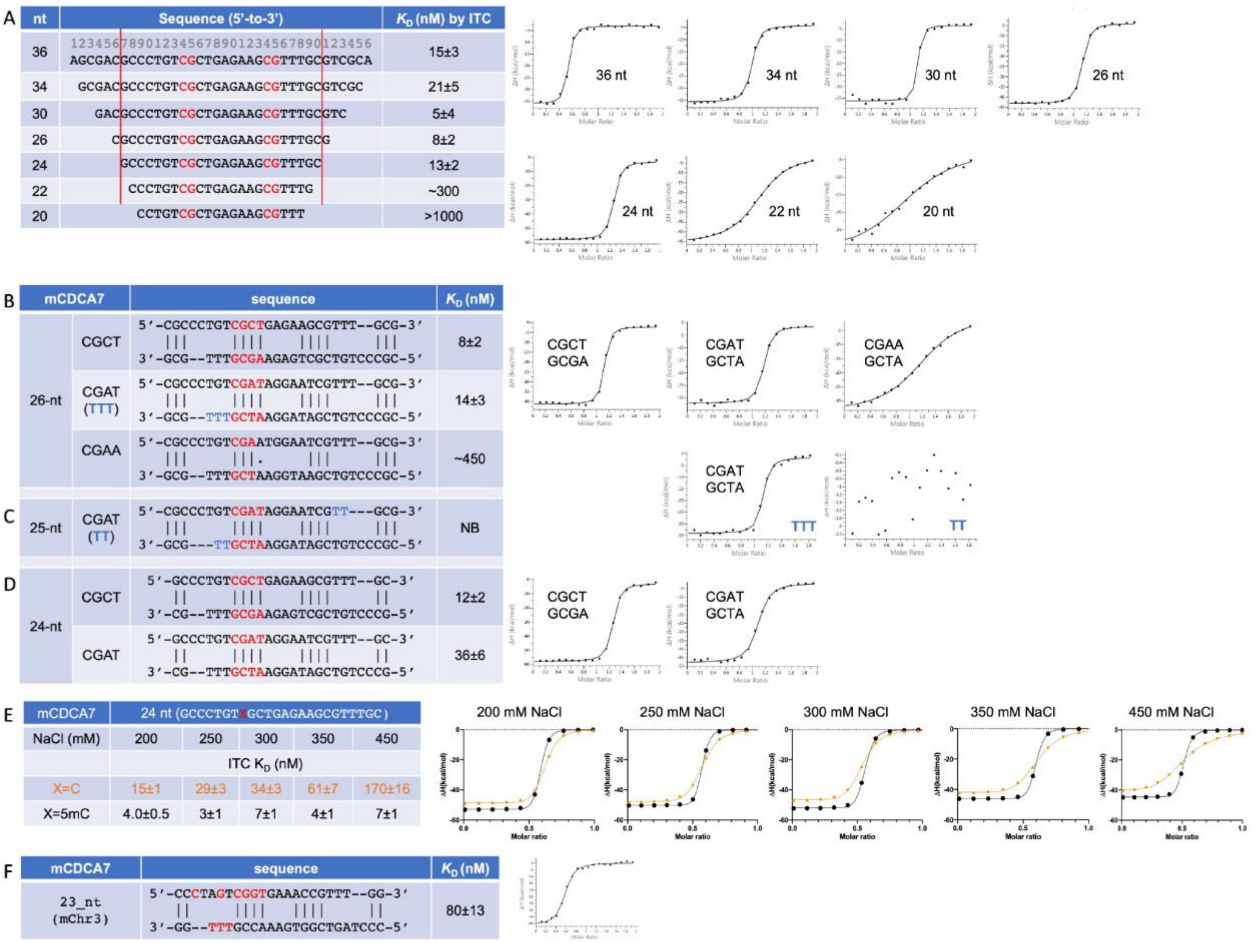


**Fig. S7. Summary of mCDCA7 CRD binding with oligonucleotides measured by ITC.** (**A**) The effect of oligonucleotide length. (**B**) The effect of the 4-bp stem S2 in the context of 26-nt. (**C**) The effect of reducing 3T-triplet to 2T. (**D**) The effect of changing CGCT to CGAT of the 4-bp stem S2 in the context of 24-nt. (**E**) The effect of ionic strength on binding with unmethylated and hemi-methylated oligos in the contact of 24-nt. (**F**) An example of mCDCA7 CRD domain binding a sequence from mouse chromosome 3.

A (hCDCA7)

<table>
<tr><th>nt</th><th>sequence</th><th>$K_D$ (nM)</th></tr>
<tr><td>26</td><td>5′-CGCCCTGTCGCTGAGAAGCGTTT--GCG-3′<br>|||   ||||   ||||   |||<br>3′-GCG--TTTGCGAAGAGTCGCTGTCCCGC-5′</td><td>14±1</td></tr>
<tr><td>24<br>(CGCT)</td><td>5′-GCCCTGTCGCTGAGAAGCGTTT--GC-3′<br>||   ||||   ||||   ||<br>3′-CG--TTTGCGAAGAGTCGCTGTCCCG-5′</td><td>13±2</td></tr>
<tr><td>22</td><td>5′-CCCTGTCGCTGAGAAGCGTTTG-3</td><td>~260</td></tr>
<tr><td>20</td><td>5′-CCTGTCGCTGAGAAGCGTTT-3′</td><td>>1000</td></tr>
<tr><td></td><td></td><td></td></tr>
<tr><td>24<br>(CGAT)</td><td>5′-GCCCTGTCGATAGGAATCGTTT--GC-3′<br>||:   ||||   ||||   ||<br>3′-CG--TTTGCTAAGGATAGCTGTCCCG-5′</td><td>35±6</td></tr>
</table>

B (hCDCA7)

<table>
<tr><th>26-nt</th><th>sequence</th><th>$K_D$ (nM)</th></tr>
<tr><td>CGCT</td><td>5′-CGCCCTGTCGCTGAGAAGCGTTT--GCG-3′<br>|||   ||||   ||||   |||<br>3′-GCG--TTTGCGAAGAGTCGCTGTCCCGC-5′</td><td>8±2</td></tr>
<tr><td>CGAT</td><td>5′-CGCCCTGTCGATAGGAATCGTTT--GCG-3′<br>|||   ||||   ||||   |||<br>3′-GCG--TTTGCTAAGGATAGCTGTCCCGC-5′</td><td>16±3</td></tr>
<tr><td>CGAA</td><td>5′-CGCCCTGTCGAATGGAATCGTTT--GCG-3′<br>|||   |||   |||   |||<br>3′-GCG--TTTGCTAAGGTAAGCTGTCCCGC-5′</td><td>~450</td></tr>
</table>

C (hCDCA7)

<table>
<tr><th>nt</th><th>sequence</th><th>$K_D$ (nM)</th></tr>
<tr><td>26<br>(TTT)</td><td>5′-CGCCCTGTCGATAGGAATCGTTT--GCG-3′<br>|||   ||||   ||||   |||<br>3′-GCG--TTTGCTAAGGATAGCTGTCCCGC-5′</td><td>12±3</td></tr>
<tr><td>26<br>(TT)</td><td>5′-CGCCCTGTCGATAGGAATCGTT---GCG-3′<br>|||   ||||   ||||   |||<br>3′-GCG---TTGCTAAGGATAGCTGTCCCGC-5′</td><td>NB</td></tr>
</table>

D (hCDCA7)

<table>
<tr><th>nt</th><th>sequence</th><th>$K_D$ (nM)</th></tr>
<tr><td>26<br>(CCTGT)</td><td>5′-CGCCCTGTCGATAGGAATCGTTT--GCG-3′<br>|||   ||||   ||||   |||<br>3′-GCG--TTTGCTAAGGATAGCTGTCCCGC-5′</td><td>23±3</td></tr>
<tr><td>26<br>(TCTAT)</td><td>5′-CGCTCTATCGATAGGAATCGTTT--GCG-3′<br>|||   ||||   ||||   |||<br>3′-GCG--TTTGCTAAGGATAGCTATCTCGC-5′</td><td>>1000</td></tr>
<tr><td>26<br>(GCTCT)</td><td>5′-CGCGCTCTCGATAGGAATCGTTT--GCG-3′<br>|||   ||||   ||||   |||<br>3′-GCG--TTTGCTAAGGATAGCTCTCGCGC-5′</td><td>NB</td></tr>
</table>

E (hCDCA7)

<table>
<tr><th>nt</th><th>sequence</th><th>$K_D$ (nM)</th></tr>
<tr><td>19<br>(2 purine<br>mismatches)</td><td>5′-GCCCTGTCGCTGAGAAGGC-3′<br>||   ||||   ||||||<br>3′-CG--TTTGCGAAGTCTCCG-5′</td><td>36±5</td></tr>
<tr><td>19<br>(3 purine<br>mismatches)</td><td>5′-GCCCTGTCGCTGAGAAGGC-3′<br>||   ||||   |||||<br>3′-CG--TTTGCGAAGATTCCG-5′</td><td>73±12</td></tr>
<tr><td>19<br>(2 purine<br>mismatches)</td><td>5′-GCCCTGTCGCTGAGAAGGC-3′<br>||   ||||   ||||<br>3′-CG--TTTGCGAAGAGTCCG-5′</td><td>~400</td></tr>
<tr><td></td><td></td><td></td></tr>
<tr><td>20<br>(7 bp 3′ end)</td><td>5′-GCCCTGTCGCTGAGAAGGGC-3′<br>||   ||||   |||||||<br>3′-CG--TTTGCGAAGCTTCCCG-5′</td><td>53±4</td></tr>
<tr><td>21<br>(8 bp 3′ end)</td><td>5′-GCCCTGTCGCTGAGAAGGGCT-3′<br>||   ||||   ||||||||<br>3′-CG--TTTGCGAAGCTTCCCGA-5′</td><td>29±6</td></tr>
<tr><td>22<br>(9 bp 3 ′ end)</td><td>5′-GCCCTGTCGCTGAGAAGGGCTA-3′<br>||   ||||   |||||||||<br>3′-CG--TTTGCGAAGCTTCCCGAT-5′</td><td>53±9</td></tr>
</table>

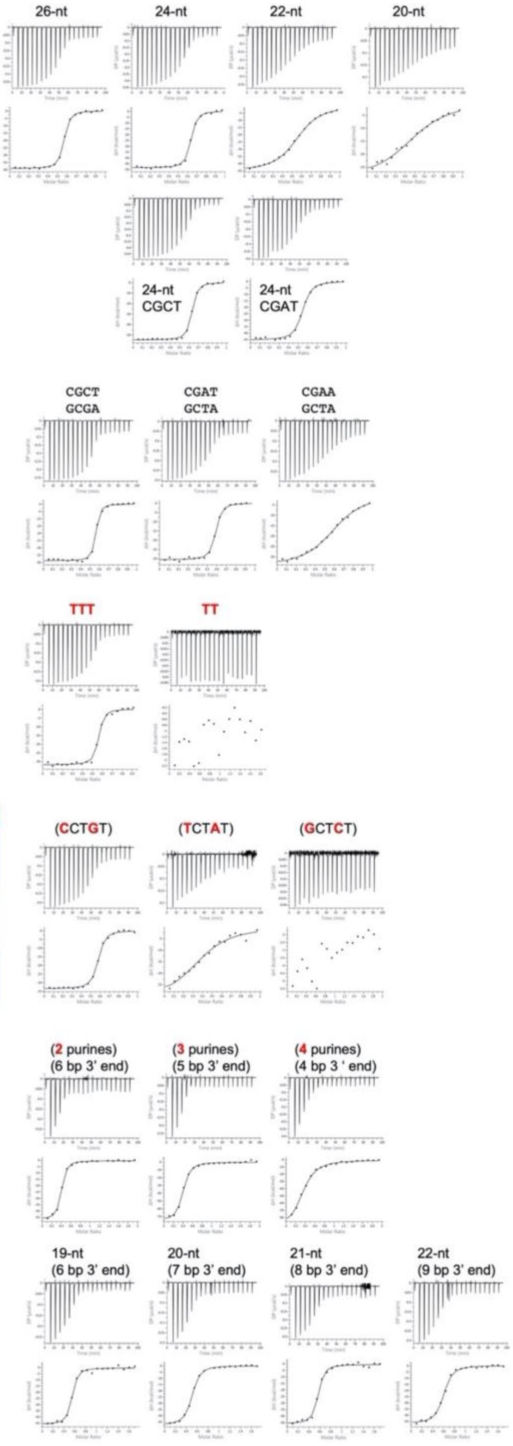


**Fig. S8. Summary of hCDCA7 CRD binding of oligonucleotides measured by ITC.** (**A**) The effect of oligonucleotide length varying from 26-nt to 20-nt. Also shown is the effect of changing CGCT to CGAT of stem S2 in the 24-nt context. (**B**) The effect of 4-bp stem S2 in the context of 26-nt. (**C**) The effect of reducing 3T-triplet to 2T. (**D**) The effect of changing intra-strand C:G base pair to T:A and G:C within the 5-nt bulge. (**E**) The effect of the number of purine mismatches after the 4-bp stem S2 and the associated number of base pairs in the 3’ end of the top strand.

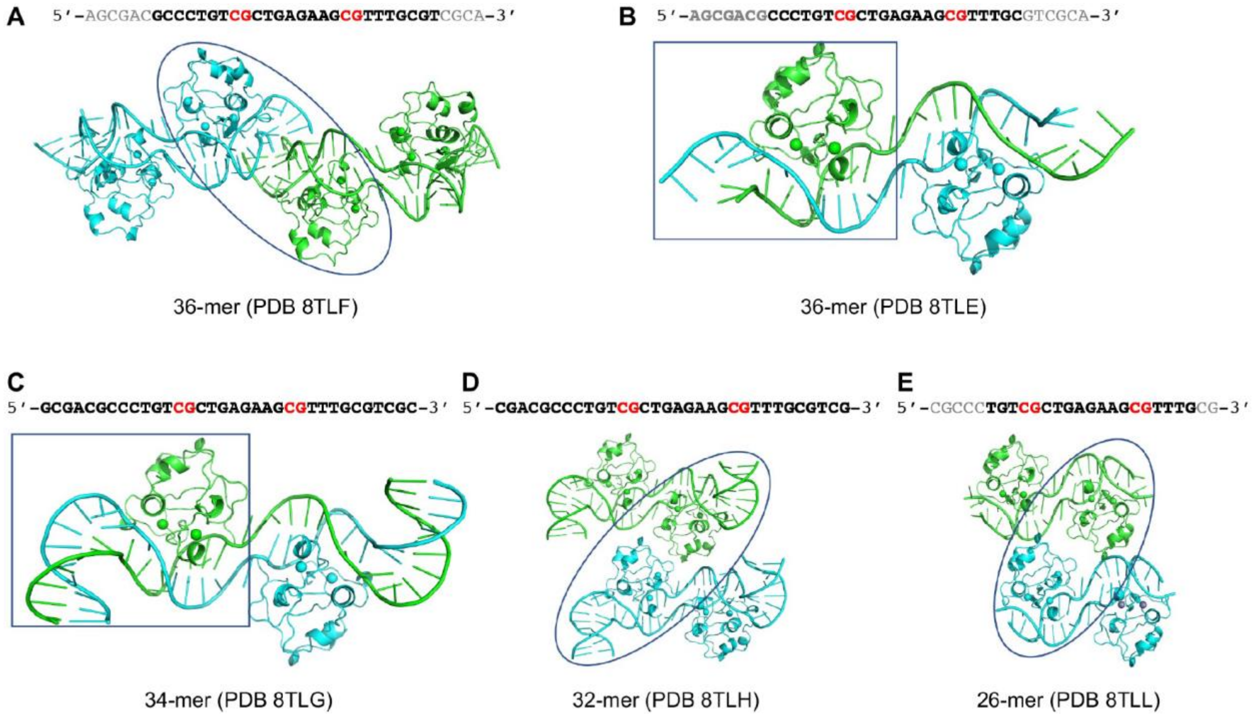


**Fig. S9. Summary of structures of mCDCA7 CRD in complex with non-B-form DNA** (for statistics, see table S1). (**A**) In the structure of PDB 8TLF, the 36-nt was used for co-crystallization, but only 26-nt (in bold) was observed in the electron density. There are two CRD-DNA complexes (circled) within the crystallographic asymmetric unit. The contact between two complexes is mainly mediated by the DNA. (**B**) In the structure of PDB 8TLE, the 23 (in bold) out of 36-nt was observed in the density. There is one CRD-DNA complex (boxed) in the crystallographic asymmetric unit. (**C**) In the structure of PDB 8TLG, the entire 34-nt was observed and there is one CRD-DNA complex per asymmetric unit. (**D**) In the structure of PDB 8TLH, the entire 32-nt was observed and there are two CRD-DNA complexes per asymmetric unit. The contact between the two complexes is mainly mediately by the protein. (**E**). In the structure of PDB 8TLL, 19 (in bold) out of 26-nt was observed and there are two CRD-DNA complexes per asymmetric unit. The contact between the two complexes is mediated by protein interaction with the neighboring DNA.

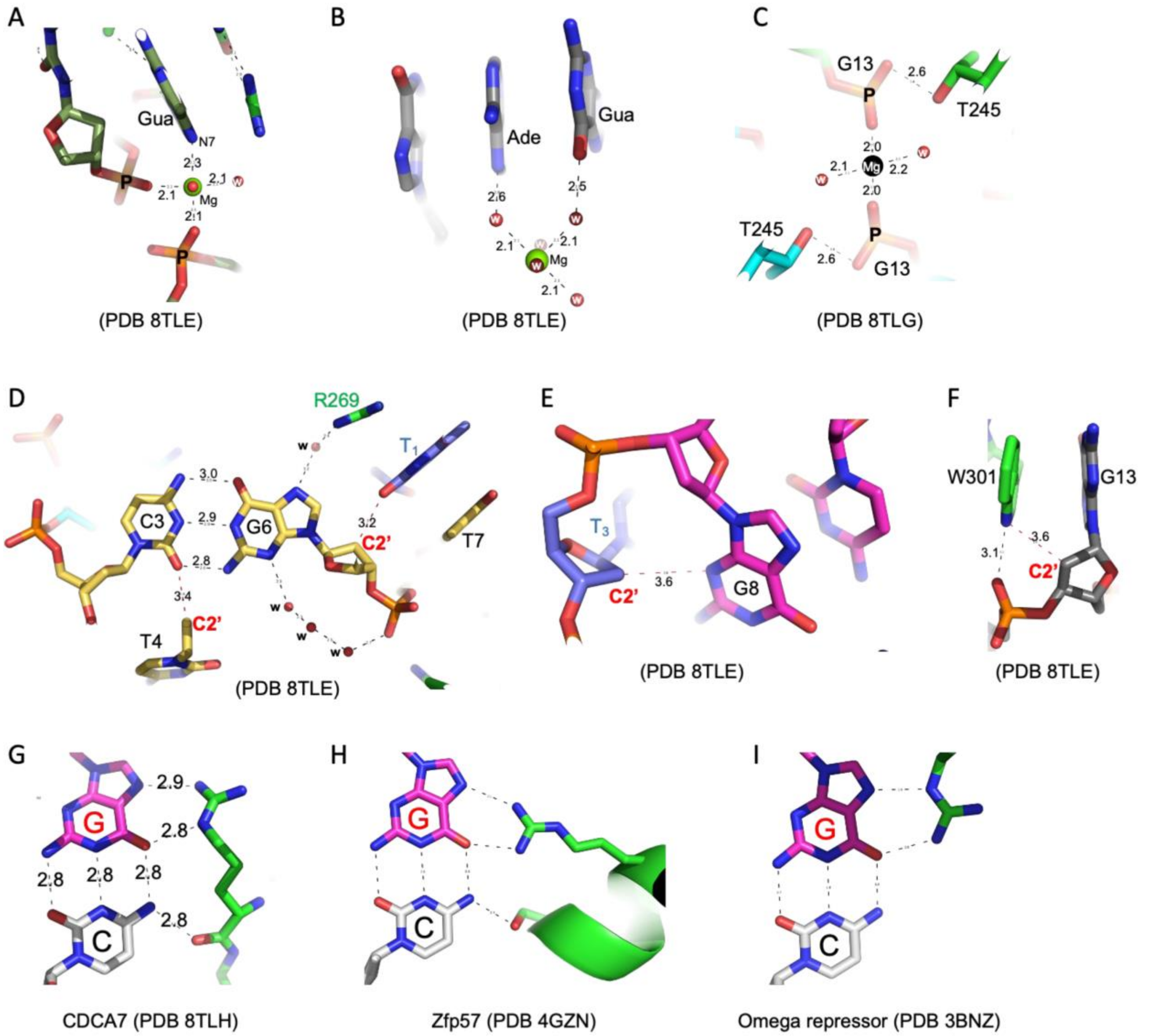


**Fig. S10. Detailed information on CDCA7-DNA interactions. (A-C) Examples of divalent $Mg^{2+}$ ions** involved in mediating DNA phosphate and base interactions (panel A), unpaired DNA bases (panel B), and two phosphate groups (panel C). The bound $Mg^{2+}$ ion is octahedrally coordinated by oxygen atoms of DNA phosphate groups and water molecules. (**D-F**) **Examples of deoxyribose *C2'* atom** (labeled in red) involved in intra-molecule or inter-molecule van der Waals contacts: *C2'* atoms of T4 and G6 (panel D), *C2'* atom of $T_3$ (panel E) and *C2'* atom of G13 (panel F). (**G-I**) **Three possible Arg conformation of forming bidentate interactions between an arginine and a guanine base.** (**G**) The side-on approach entry from the cytosine side of the C:G base pair. The Arg interacts with both bases and cytosine methylation would interfere with the close contact between Arg-Gua. (**H**) The end-on approach using both of the distal nitrogen atoms. (**I**) The side-on approach entry from the guanine side. In panels H and I, the Arg interacts only with the guanine base, and cytosine methylation could be accommodated.

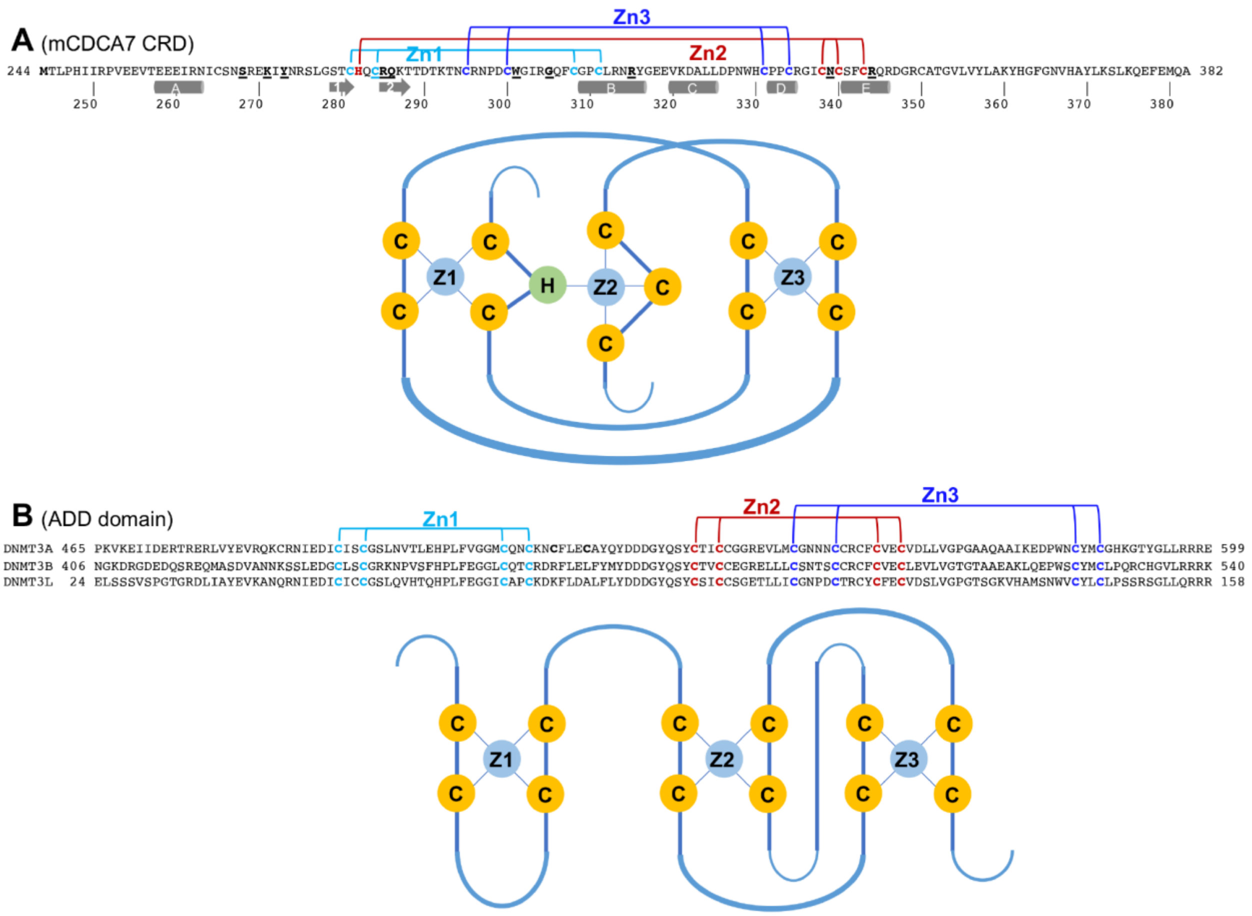


**Fig. S11. Comparison of zinc coordination.** (**A**) CDCA7 CRD has a unique cross-braced zinc coordination by CHQC, where the two Cys residues and His residue contribute to binding of two different zinc ions. (**B**) The ADD domain of ATRX, DNMT3A/3B and DNMT3L consists of an N-terminal GATA-like zinc finger and a plant homeodomain finger (PHD) which binds histone tail peptides.

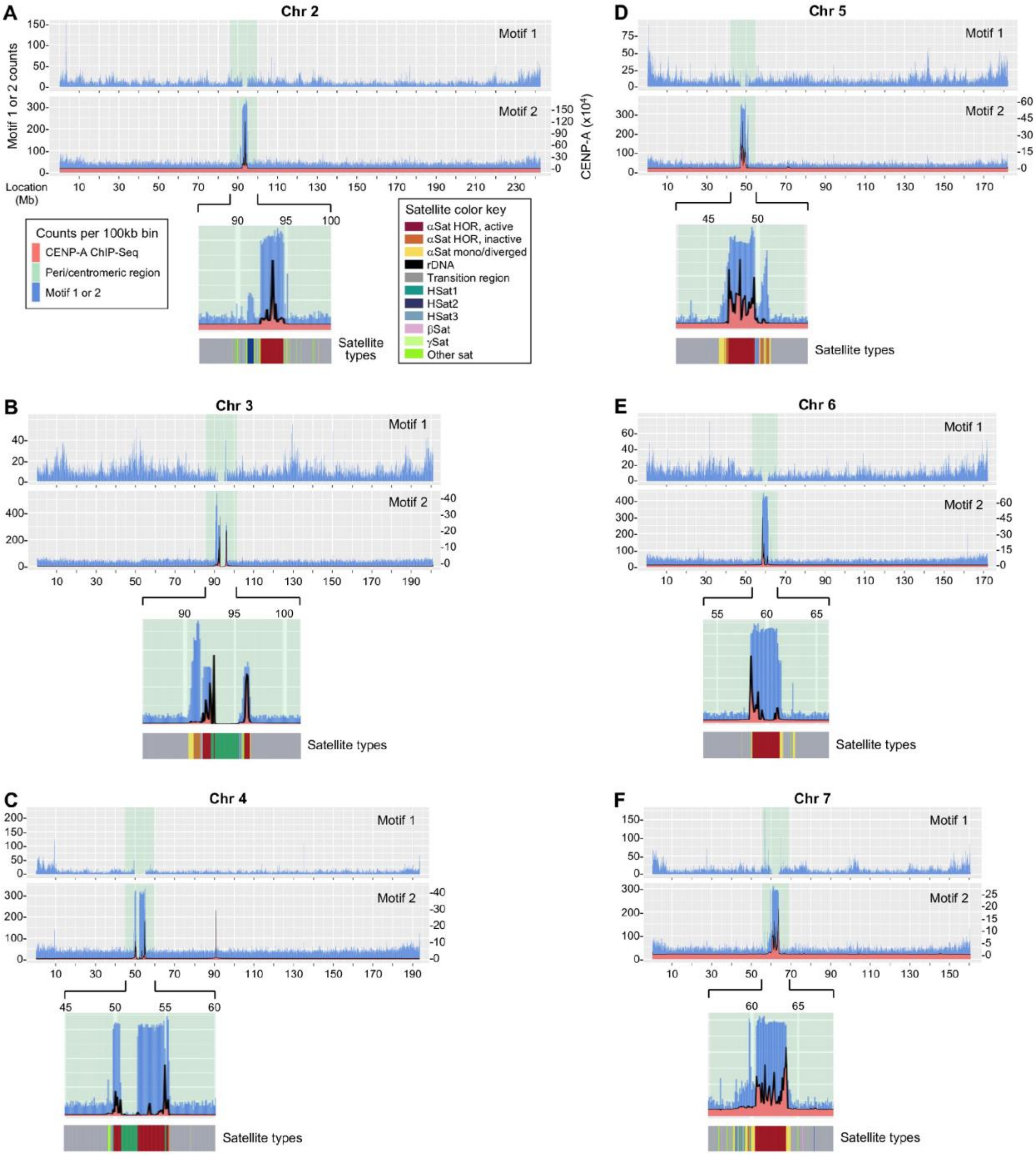
A
Chr 2
Motif 1
Motif 2
Motif 1 or 2 counts
CENP-A (x10⁴)
Location (Mb)
Counts per 100kb bin
CENP-A ChIP-Seq
Peri/centromeric region
Motif 1 or 2
Satellite color key
αSat HOR, active
αSat HOR, inactive
αSat mono/diverged
rDNA
Transition region
HSat1
HSat2
HSat3
βSat
γSat
Other sat
Satellite types
B
Chr 3
C
Chr 4
D
Chr 5
E
Chr 6
F
Chr 7

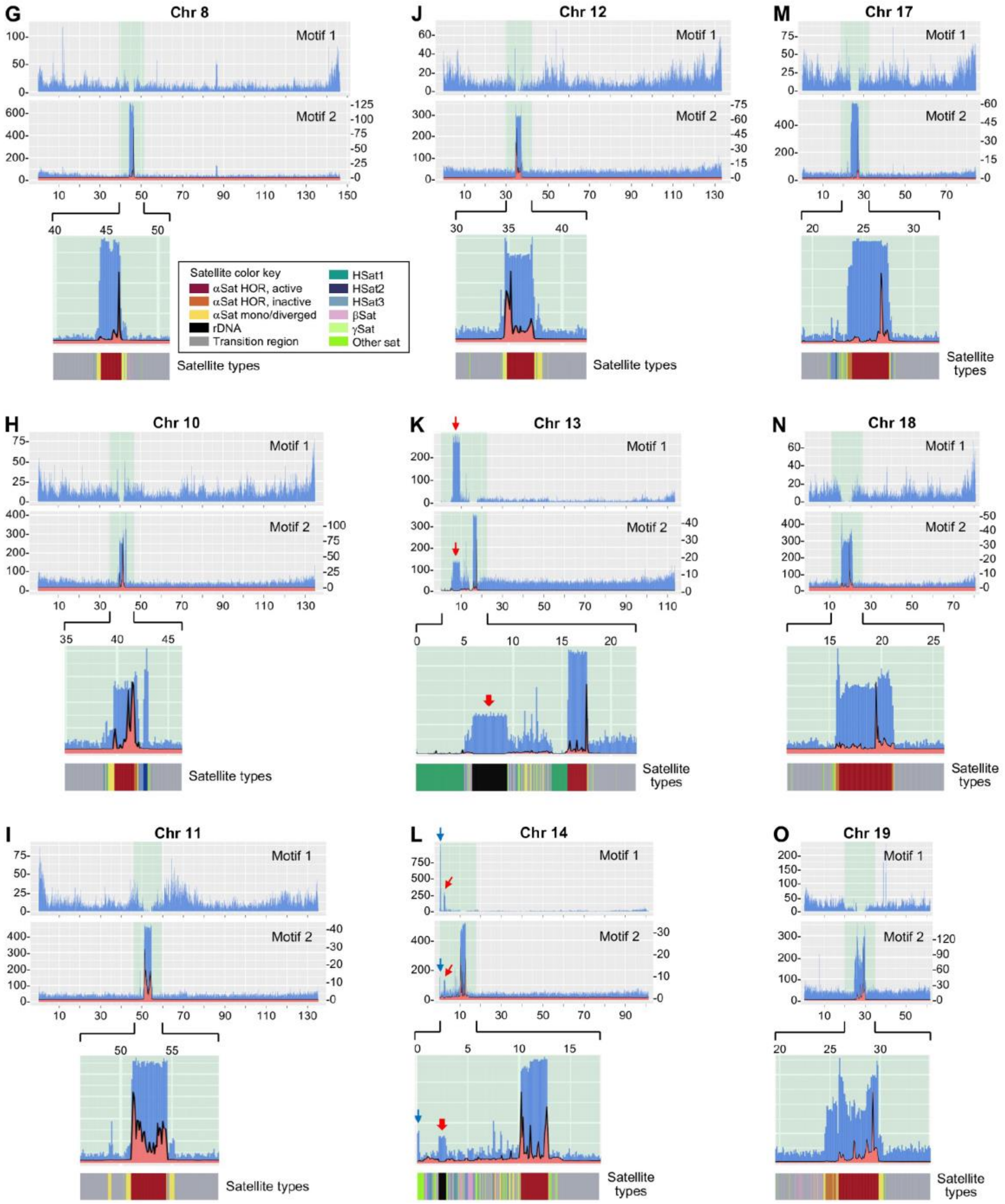

G
Chr 8
J
Chr 12
M
Chr 17
H
Chr 10
K
Chr 13
N
Chr 18
I
Chr 11
L
Chr 14
O
Chr 19
Motif 1
Motif 2
Satellite types
Satellite color key
αSat HOR, active
αSat HOR, inactive
αSat mono/diverged
rDNA
Transition region
HSat1
HSat2
HSat3
βSat
γSat
Other sat

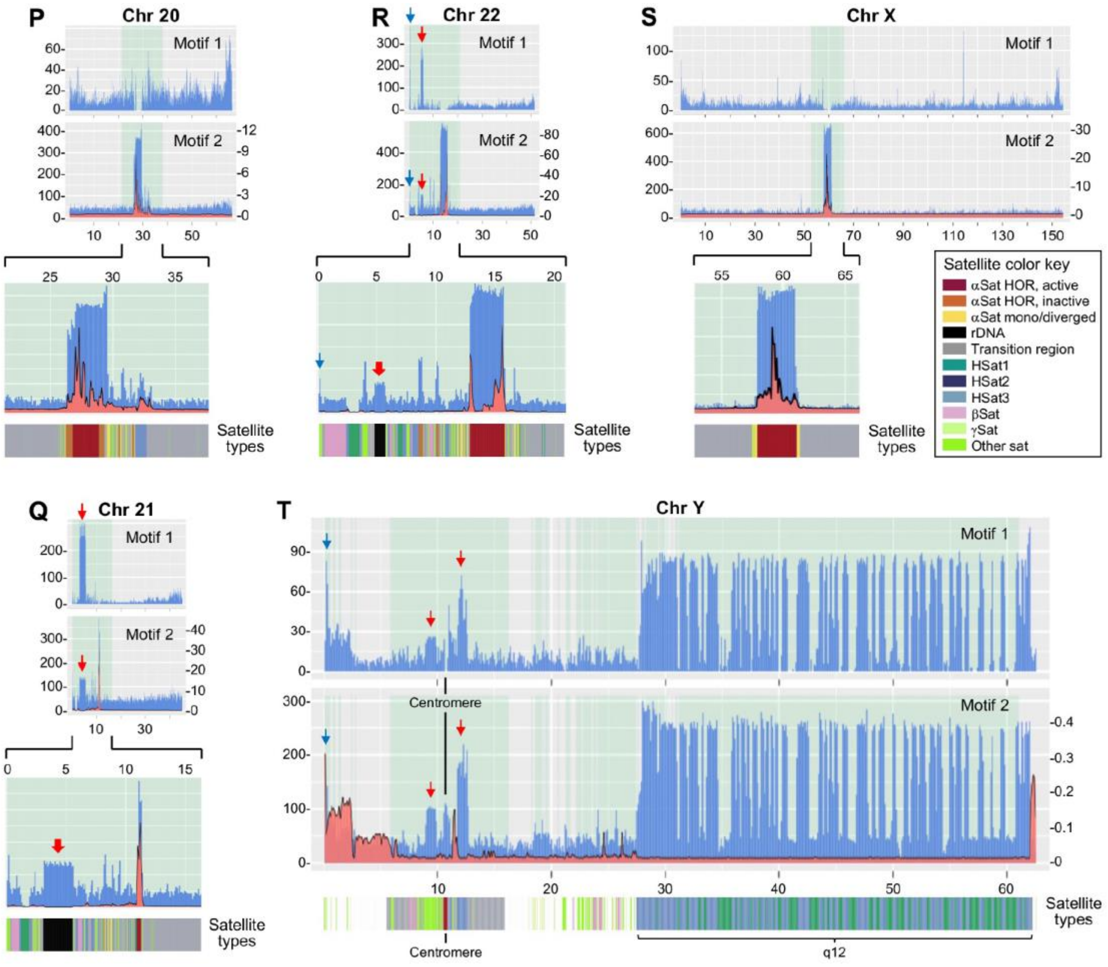


**Fig. S12. Distribution and abundance of motifs 1 and 2 on human chromosomes.** Shown are results of all chromosomes except chr 1, 9, 15 and 16 (which are presented in Fig. 5). The red and blue arrows indicate motifs 1 and 2 peaks at the same locations in the pericentromeric regions (red) or at the end of p arms (blue) of the acrocentric chromosomes (chr 13, 14, 15, 21 and 22) and chr Y (**K, L, Q, R**, **T** and Fig. 5C for chr 15). Unlike the other chromosomes, which are shown proportionally in sizes, with the peri/centromeric regions being expanded 5-fold, the entire Y chromosome is shown at a larger magnification without expanding the peri/centromeric region, and the centromere and the large heterochromatic region at the distal end of the q arm (q12) are indicated (**T**).

### Table S1. Summary of X-ray data collection and refinement statistics (*)

| Protein Construct | mouse CDCA7 | mouse CDCA7 | mouse CDCA7 | mouse CDCA7 | mouse CDCA7 | human CDCA7 | mouse CDCA7 |
|---|---|---|---|---|---|---|---|
| DNA | 36-nt | 36-nt | 34-nt | 32-nt | 32-nt (5mC) | 32-nt (5mC) | 26-nt |
| PDB Code | **8TLE** | **8TLF** | **8TLG** | **8TLH** | **8TLJ** | **8TLK** | **8TLL** |
| Date Collected | 03/10/2020 | 02/06/2020 | 07/14/2020 | 09/25/2020 | 05/17/2021 | 05/17/2021 | 09/25/2020 |
| Beamline | APS, 22ID | APS, 22ID | APS, 22ID | APS, 22ID | ALS, 502 | ALS, 502 | APS, 22ID |
| Wavelength (Å) | 1.00000 | 1.00000 | 1.00000 | 1.00000 | 0.99999 | 0.99999 | 1.00000 |
| Space group | *C*2 | *C*2 | *C*2 | *C*2 | *C*2 | *C*2 | *C*2 |
| Cell dimensions (Å) | 88.67, 60.77, 49.09 | 89.40, 31.49, 133.3 | 86.49, 60.56, 53.23 | 169.68, 31.52, 91.63 | 169.87, 31.53, 91.82 | 173.48, 31.54, 94.57 | 87.71, 60.72, 84.11 |
| α, β, γ (°) | 90, 117.5, 90 | 90, 95.75, 90 | 90, 123.10, 90.0 | 90, 110.59, 90 | 90, 110.8, 90 | 90, 110.2, 90 | 90, 114.0, 90 |
| Resolution (Å) | 30.38-2.08 (2.18-2.08) | 29.90-1.64 (1.71-1.64) | 30.28-2.09 (2.16-2.09) | 39.71-1.94 (2.02-1.94) | 45.03-2.30 (2.37-2.30) | 43.06-2.99 (3.09-2.99) | 48.40-2.59 (2.68-2.58) |
| [a] $R_{merge}$ | 0.165 (0.490) | 0.095 (0.509) | 0.068 (0.342) | 0.080 (0.678) | 0.061 (0.419) | 0.123 (0.449) | 0.199 (0.609) |
| $R_{pim}$ | 0.047 (0.178) | 0.049 (0.343) | 0.037 (0.257) | 0.035 (0.464) | 0.038 (0.321) | 0.070 (0.333) | 0.072 (0.444) |
| $CC_{1/2}$ | 0.990 (0.919) | 0.992 (0.807) | 1.006 (0.827) | 0.997 (0.723) | 0.997 (0.832) | 1.000 (0.777) | 0.967 (0.702) |
| [b] $\langle I/\sigma I \rangle$ | 13.1 (3.7) | 13.6 (2.2) | 15.5 (2.8) | 17.8 (1.1) | 16.7 (1.6) | 6.1 (1.7) | 8.7 (1.6) |
| Completeness (%) | 98.8 (94.0) | 89.9 (65.4) | 83.6 (47.7) | 84.8 (34.4) | 95.1 (69.0) | 79.7 (75.8) | 93.6 (63.3) |
| Redundancy | 11.3 (5.9) | 3.9 (2.1) | 3.6 (2.1) | 5.2 (2.1) | 3.1 (1.9) | 3.6 (2.3) | 7.0 (1.9) |
| Observed reflections | 148, 936 | 158,005 | 41,440 | 151,288 | 62,043 | 29,031 | 83,535 |
| Unique reflections | 13,164 (641) (12,082 have I+ and I-) | 40,904 (2,920) | 11,460 (653) | 29,139 (1,162) | 19,752 (1,408) | 8135 (772) | 12,017 (814) |
| Mean FOM (SAD) | 0.314 (at 4 Å) | | | | | | |
| Density Modification R-factor | 0.42 (at 4 Å) | | | | | | |
| **Refinement** | | | | | | | |
| Resolution (Å) | 2.08 | 1.64 | 2.09 | 1.94 | 2.30 | 2.99 | 2.58 |
| No. reflections | 13,033 | 40,879 | 11,449 | 29,047 | 19,726 | 8002 | 11,979 |
| [c] $R_{work}$ / [d] $R_{free}$ | 0.227 / 0.262 | 0.188 / 0.212 | 0.180 / 0.219 | 0.200 / 0.223 | 0.183 / 0.228 | 0.238 / 0.298 | 0.242 / 0.285 |
| No. Atoms | (1 complex) | (2 complexes) | (1 complex) | (2 complexs) | (2 complexes) | (2 complexes) | (2 complxes) |
| Protein | 832 | 1683 | 817 | 1630 | 1630 | 1638 | 1578 |
| DNA | 470 | 1052 | 694 | 1306 | 1314 | 1308 | 767 |
| Zn | 3 | 6 | 3 | 6 | 6 | 6 | 6 |
| Solvent | 15 | 283 | 75 | 160 | 140 | 87 | 24 |
| B Factors ($Å^2$) | | | | | | | |
| Protein | 85.4 | 31.5 | 48.1 | 56.0 | 58.1 | 32.7 | 54.1 |
| DNA | 96.8 | 26.8 | 79.0 | 80.3 | 79.7 | 53.1 | 58.3 |
| Zn | 62.5 | 18.8 | 36.4 | 43.9 | 47.1 | 20.2 | 44.8 |
| Solvent | 85.9 | 32.9 | 46.8 | 53.5 | 53.4 | 26.9 | 48.1 |
| **R.m.s. deviations** | | | | | | | |
| Bond lengths (Å) | 0.003 | 0.002 | 0.004 | 0.003 | 0.004 | 0.003 | 0.003 |
| Bond angles (°) | 0.5 | 0.5 | 0.7 | 0.5 | 0.7 | 0.5 | 0.5 |

* Values in parenthesis correspond to highest resolution shell.

[a] $R_{merge} = \Sigma \mid I - \langle I \rangle \mid / \Sigma I$, where I is the observed intensity and <I> is the averaged intensity from multiple observations.

[b] $\langle I/\sigma I \rangle$ = averaged ratio of the intensity (I) to the error of the intensity (σI).

[c] $R_{work} = \Sigma \mid Fobs - Fcal \mid / \Sigma \mid Fobs \mid$, where Fobs and Fcal are the observed and calculated structure factors, respectively.

[d] $R_{free}$ was calculated using a randomly chosen subset (5%) of the reflections not used in refinement.

### Table S5. Primers and synthesized DNA

| Name | Sequence (5' to 3') | Notes |
|---|---|---|
| C-2646 | GTTCTAGCGGCCGCTAGAAGAGTGCCTCAGAAGGATCTCAGAGTGAAGAAG<br>AACCTGAAGAAGTTCAGATATGTGAAGTTGATCTCCATGGAGACCTCGTCA<br>TCCTCTGATGACAGTTGTGACAGCTTCGCTTCTGATAACTTCGCCAACACG<br>AGGCTGCAGTCAGTTAGAGAAGGCTGTAGGACACGCAGCCAGTGCAGGCAC<br>TCTGGACCTCTCAGGGTGGCGATGAAGTTTCCAGCGCGGAGTACCAGAGGA<br>GCAACCAACAAGAAAGCAGAGTCTAGACAGCCCTCAGAGAACTCTGTGACT<br>GATTCCAACTCCGATTCAGAAGATGAGAGTGGAATGAACTTCTTGGAGAAG<br>AGAGCCTTGAATATCAAGCAGAACAAGGCAATGCTTGCCAAGCTCATGTCT<br>GAATTAGAGAGCTTCCCTGGCTCGTTCCGTGGAAGACATCCTCTCCCAGGC<br>TCCGACTCACAATCAAGGAGACCTCGAAGGCGTACATTCCCAGGTGTTGCT<br>TCCAGGAGAAACCCTGAACGGAGAGCTCGTCCTCTTACCAGGTCAAGGTCC<br>CGGATCCTCGGGTCCCTTGACGCTCTACCTATGGAGGAAGAAGAGGAAGAG<br>GATAAGTACATGTTGGTGAGGAAGAGGAAGACCGTGGATGGCTACATGAAT<br>GAAGATGACCTGCCCAGAAGCCGTCGCTCCAGATCATCCGTGACCCTTCCG<br>CATATAATTCGCCCAGTGGAAGAGATTACAGAGGAAGAGTTGGAGAACGTC<br>TGCAGCAATTCTCGAGAGAAGATATATAACCGTTCACTGGGCTCTACTTGT<br>CATCAATGCCGTCAGAAGACTATTGATACCAAGACCAACTGCAGGAACCCA<br>GACTGCTGGGGAGTTCGAGGCCAGTTCTGTGGACCTTGCCTTCGGAACCGT<br>TATGGTGAAGAGGTCAGAGATGCTCTGCTGGACCCGAACTGGCATTGTCCA<br>CCTTGTCGAGGAATCTGCAACTGCAGCTTCTGCAGACAGCGAGATGGACGG<br>TGTGCGACTGGAGTCCTTGTGTATTTAGCCAAGTATCATGGCTTCGGAAT<br>GTGCATGCCTACTTGAAGAGCCTGAAGCAGGAATTTGAAATGCAAGCATGA<br>ATTCAGACTG | Synthesized *hCDCA7* cDNA for HA-hCDCA7 construct. Some codons were changed to synonymous ones. |
| C-1257 (F)<br>C-1258 (R) | CCAGCGGCCGCCGAACAAACGGAGCCTGC<br>CTTGCGGCCGCATTAAAATAAACATTCAGC | Amplify *mHells* cDNA for HA-mHELLS construct |
| C-1447 (F)<br>C-1248 (R) | AACGAATTCGCTCGCCGCGCGCGGCAAAAG<br>TTCGAATTCTACGCTTGCATTTCAAATTC | Amplify *mCdca7* cDNA for GFP-mCDCA7 construct |
| C-1448 (F)<br>C-1248 (R) | AGTGAATTCCCTGGATCCATGACCCTTCCG<br>TTCGAATTCTACGCTTGCATTTCAAATTC | Amplify *mCdca7* cDNA fragment for GST-mCDCA7 CRD construct |
| C-2142 (F)<br>C-2642 (R) | TCCGAATTCCTTCCGCATATAATTCGC<br>TCTGAATTCATGCTTGCATTTC | Amplify *hCDCA7* cDNA fragment for GST-hCDCA7 CRD construct |
| C-1412 (F)<br>C-1411 (R) | CATCAGTGTC**A**CCAGAAAACCACTGACAC<br>GGTTTTCTGG**T**GACACTGATGACATGTAG | Introduce the R285H mutation into mCDCA7 |
| C-1498 (F)<br>C-1497 (R) | GGCATCCGGG**T**CCAATTCTGTGGTCCCT<br>ACAGAATTGG**A**CCCGGATGCCCCAGCAGTC | Introduce the G305V mutation into mCDCA7 |
| C-1500 (F)<br>C-1499 (R) | CTTCGAAACC**A**CTATGGCGAGGAGGTCAAG<br>CTCGCCATAG**T**GGTTTCGAAGGCAGGGAC | Introduce the R315H mutation into mCDCA7 |
| C-1496 (F)<br>C-1495 (R) | CATGAACTTT[ ]GAAAGCTTCCCTGGCTTG<br>GGAAGCTTTC[ ]AAAGTTCATGCCATCTTC | Delete 66 nt in *mCdca7* cDNA to generate mCDCA7ΔLZ |
| C-1447 (F)<br>C-1615 (R) | AACGAATTCGCTCGCCGCGCGCGGCAAAAG<br>ACTGAATTCATGTAGAACCCAGAGAGC | Amplify *mCdca7* cDNA fragment for GFP-mCDCA7ΔCRD construct |
| C-2145 (F)<br>C-2144 (R) | CATCAATGCC**A**TCAGAAGACTATTGATAC<br>AGTCTTCTGA**T**GGCATTGATGACAAGTAG | Introduce the R274H mutation into hCDCA7 |
| SELEX-F<br>SELEX-R | TAGGGAAGAGAAGGACATATGAT<br>TCAAGTGGTCATGTACTAGTCAA | Amplify bound DNA to generate ssDNA pool after each round of SELEX |

Underlined are restriction sites used for cloning; shown in bold are mutations; [ ] indicates location of deletion

**Table S2. (separate Excel file)**
Paired motifs 1 and 2 in peaks on acrocentric chromosomes.

**Table S3. (separate Excel file)**
Motif 1 counts in peri/centromeric regions.

**Table S4. (separate Excel file)**
Motif 2 counts in peri/centromeric regions.
